# FEMA: Fast and efficient mixed-effects algorithm for large sample whole-brain imaging data

**DOI:** 10.1101/2021.10.27.466202

**Authors:** Pravesh Parekh, Chun Chieh Fan, Oleksandr Frei, Clare E. Palmer, Diana M. Smith, Carolina Makowski, John R. Iversen, Diliana Pecheva, Dominic Holland, Robert Loughnan, Pierre Nedelec, Wesley K. Thompson, Donald J. Hagler, Ole A. Andreassen, Terry L. Jernigan, Thomas E. Nichols, Anders M. Dale

## Abstract

The linear mixed-effects model (LME) is a versatile approach to account for dependence among observations. Many large-scale neuroimaging datasets with complex designs have increased the need for LME, however LME has seldom been used in whole-brain imaging analyses due to its heavy computational requirements. In this paper, we introduce a fast and efficient mixed-effects algorithm (FEMA) that makes whole-brain vertex-wise, voxel-wise, and connectome-wide LME analyses in large samples possible. We validate FEMA with extensive simulations, showing that the estimates of the fixed effects are equivalent to standard maximum likelihood estimates but obtained with orders of magnitude improvement in computational speed. We demonstrate the applicability of FEMA by studying the cross-sectional and longitudinal effects of age on region-of-interest level and vertex-wise cortical thickness, as well as connectome-wide functional connectivity values derived from resting state functional MRI, using longitudinal imaging data from the Adolescent Brain Cognitive Development^SM^ Study release 4.0. Our analyses reveal distinct spatial patterns for the annualized changes in vertex-wise cortical thickness and connectome-wide connectivity values in early adolescence, highlighting a critical time of brain maturation. The simulations and application to real data show that FEMA enables advanced investigation of the relationships between large numbers of neuroimaging metrics and variables of interest while considering complex study designs, including repeated measures and family structures, in a fast and efficient manner. The source code for FEMA is available via: https://github.com/cmig-research-group/cmig_tools/.

## 1. Introduction

As neuroimaging studies have moved to large sample sizes, the size of datasets and the complexity of relationships among observations within these datasets pose several challenges to neuroimaging analysis. For example, in the Adolescent Brain Cognitive Development^SM^ Study (ABCD Study^®^) [https://abcdstudy.org], the inclusion of twins and siblings, in addition to a longitudinal design, make it necessary to simultaneously consider multiple correlations among observations. Similarly, other large-scale studies such as the UK Biobank include repeated observations and other structures in the data that introduce correlations among observations within the dataset [https://www.ukbiobank.ac.uk]. Mixed-effects models are a flexible method of modeling such known patterns in the data. Within neuroimaging, linear mixed-effects models (LME) have previously been used to model repeated measurements, family relatedness, and longitudinal observations (see, for example, (Beck et al., 2021; Bernal-Rusiel et al., 2013; Chen et al., 2013; Dick et al., 2021; Fjell et al., 2021; Nyberg et al., 2021; Stoffers et al., 2015; Varosanec et al., 2015)).

Within an LME, study design and heterogeneity in the study population can be parameterized by specifying the covariance pattern for the random effects; accounting for the actual correlations in the data by means of random effects can lead to an increase in statistical power and reduction of inferential biases (Bernal-Rusiel et al., 2013). These statistical characteristics make the LME an increasingly important tool in analyzing large-scale data with known structures. However, there has been limited use of LME in brain imaging studies, especially when considering imaging analyses (e.g., vertex-wise, voxel-wise, and connectome-wide). This is due to high computational demands of LMEs and the relative lack of available methods that support performing LME estimation on whole-brain neuroimaging data. Hence, the full potential of the massive large-scale imaging samples currently collected, e.g., to reveal more of the complexity of human neurodevelopment, is yet to be realized.

The estimation of model parameters in LME is typically achieved through the maximization of the likelihood or the restricted maximum likelihood (REML). These iterative methods can easily become computationally prohibitive in the case of neuroimaging analyses, especially when considering whole-brain voxel-wise, vertex-wise, or connectome-wide analyses. This is because an LME method applied to, for example, voxel-wise data will require repeated inversions or solving for each voxel; this contrasts with ordinary least squares (OLS), where a single inversion or solution can be found to estimate the coefficients for all voxels simultaneously. Therefore, the computational efficiency of estimating many voxels using OLS is profound due to efficient implementation of matrix operations, while voxel-wise covariance matrices demand a much slower approach using loops. Consider, for example, that fitting a simple LME model takes 0.5 seconds. A typical 1.5mm isotropic standard-space image has about 558,718 analyzable voxels. Therefore, fitting this model (serially) for all the voxels would take about 3 days of computational time. If the same model would have taken 30 seconds to fit per voxel (which is reasonable for large samples with multiple random effects with many levels), the total computational time would exceed 190 days (this timing can be proportionally reduced with parallel computing, but the computational burden remains large). Thus, estimating model parameters of these LMEs can be computationally prohibitive, to the extent that it may be impossible to do some of these analyses on standard computational facilities. Therefore, there is a need for a fast and efficient LME algorithm that is capable of handling large-scale imaging data. The release of data from the ABCD Study^®^ (Casey et al., 2018) makes this unmet need more salient, as best practices for statistical analysis are to control for cohort heterogeneity and relatedness due to the inclusion of multiple siblings and twins in the sample, in order to correctly model cross-sectional and longitudinal measures (Dick et al., 2021; Smith and Nichols, 2018).

As mentioned before, LMEs have been used for the analysis of neuroimaging data. For example, the analysis of functional magnetic resonance imaging (fMRI) data using Statistical Parametric Mapping (SPM) relies on a mixed-effects framework, either via a two-stage procedure (Holmes and Friston, 1998) or a full mixed model specification (Friston et al., 2005). Another example is the LME toolbox for longitudinal analyses released with Freesurfer (Bernal-Rusiel et al., 2013) which performs cortical surface vertex-wise LME by segmenting vertices into a smaller set of regions of interest (ROIs) that share similar random effects and are geodesically close, thus reducing the number of outcome variables and the size of the covariance matrix. However, it currently lacks support for complex study design such as inclusion of family members, and for imaging measures beyond cortical surfaces. A different example is the *3dLME* tool (Chen et al., 2013) implemented in AFNI which uses the *lme4* and *nlme* packages in R along with parallel computing to perform mixed-effects analyses. An alternate approach to making mixed-effects modeling practical for neuroimaging is to fit only a marginal model and rely on a sandwich estimator of standard errors to incorporate repeated measures covariances. The SwE toolbox takes this approach, initially using OLS for parameter estimation, followed by estimating intra-block covariances when conducting inference (Guillaume et al., 2014). More recently, the “Big” Linear Mixed Models (BLMM) tool has been developed (Maullin-Sapey and Nichols, 2022) that relies on massive parallelization and high-performance computing along with a novel Fisher scoring procedure (Maullin-Sapey and Nichols, 2021) for solving LMEs. We additionally refer to (Maullin-Sapey and Nichols, 2022) for a comparative note on the software available for fitting mixed models using neuroimaging data, as well as details on the challenges when using mixed models for neuroimaging data.

In this paper, we present a fast and resource-efficient LME algorithm, called the Fast and Efficient Mixed-effects Algorithm (FEMA), that can perform whole-brain image-wise analyses on complex large-scale imaging data in a computationally efficient manner. By using a method of moments estimator, efficient binning of random effects variance parameters, leveraging the sparsity of the random effects design matrices, and vectorized operations, FEMA can perform whole-brain voxel- or vertex-wise LME analyses within minutes, compared to days or even months required using standard LME solvers. In addition, by using the sparsity in the random effects design matrices, FEMA also achieves memory efficiency. These features allow FEMA to be run on an ordinary computer without the need for very large memory capacity or extensive parallelization. The goal of this manuscript is to introduce FEMA, discuss key aspects of the algorithm that make it computationally efficient, and additionally document the implementation of FEMA that will serve as a reference to readers. Having described these in detail, we will demonstrate, using realistically simulated data, that the results from FEMA are consistent with classical maximum likelihood (ML) at a substantially reduced computational burden. Finally, we will showcase the use of FEMA by applying it to the longitudinal multimodal brain imaging data from the ABCD Study^®^ to uncover neurodevelopmental patterns of human brain change during early adolescence in cortical thickness as well as resting state functional connectivity.

## 2. Materials and Methods

In this section, we will introduce the framework of mixed-effects models as implemented in FEMA. First, we present an overview of the approach. Then, we will introduce the model and the notation which we will use throughout this paper. Next, we will go into the details of the FEMA approach to estimating the mixed model parameters. Specifically, we will present how the coefficients for the fixed effects and the variance components for the random coefficients are estimated (details on the implementation in FEMA are mentioned in the *Implementation details* section of the Supplementary Materials). Finally, at the end of this section, we will present realistic simulation scenarios for testing FEMA as well as the empirical application of FEMA using data from the ABCD Study^®^.

### 2.1 Outline

The estimation of fixed effects and variance components of the random effects in FEMA follows the following steps: first, we estimate the fixed effects using an OLS solution. Then, we compute the squares and products of residuals for pairs of observations and write it out as a vector (with some sparsity); this vector is treated as the outcome variable with the random effects as the predictor variables. We employ a method of moments (MoM) estimator to obtain an estimate of the variances associated with these random effects (equivalent to an OLS solution with an additional non-negativity constraint). Having estimated the variance components, we re-estimate the fixed effects by implementing a generalized least squares (GLS) solution, resulting in an updated estimate of the fixed effects. In case of multiple outcome variables, we employ a binning strategy prior to implementing the GLS – for all outcome variables that have similar random effects, the same covariance matrix is used for implementing the GLS. The GLS estimator for fixed effects is unbiased and consistent, and the MoM approach for the estimation of varainces of the random effects is also consistent. Further, the GLS estimator is more efficient (i.e., has smaller variance) than the OLS estimator. Additional on the theoretical background and statistical properties of different estimators (ML, OLS, and GLS) can be found in (Demidenko, 2013, chap. 3; Fitzmaurice et al., 2011, pp. 92– 94; Zou et al., 2017). In the next sections, we describe each aspect of FEMA in detail.

### 2.2 Model Set Up

Let a dataset contain *N* observations indexed by *i* ∈ 1…*N* and *J* imaging measures indexed by *j* ∈ 1…*J*. Let *X* denote a matrix of *p* covariates (or fixed effects) where the columns of the matrix are individual covariates and entry *X*_*i*_ refers to the value(s) of the covariate(s) for *i*^th^ observation. Let *y* denote the outcome variables, with entry *y*_*ij*_ indicating the value of the *i*^th^ observation for the *j*^th^ imaging measure. Then, the additive linear relationship between the outcome variables, *y*, and fixed effects, *X*, can be written as:

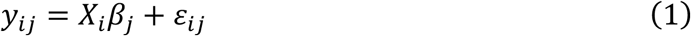

where *β*_*j*_ represents the weights associated with the fixed effects for the *j*^th^ imaging measure and *ε*_*ij*_ represents random variations for the *i*^th^ observation for the *j*^th^ imaging measure. We follow a mass univariate approach where we will not attempt to model the dependence between the *J* imaging measures, but we will combine information across the measures to obtain reliable variance parameter estimates.

In the fixed effects only setting (e.g., general linear model), *ε*_*ij*_ accounts for measurement errors for each imaging variable, uncorrelated across observations *i*. For a marginal model (e.g., generalized estimating equations), arbitrary dependence is captured in some combination of a working covariance and a robust “sandwich” standard error. The LME, on the other hand, explicitly models both the uncorrelated measurement error and correlations among observations attributable to the known structure in the data (“random effects”). For every imaging measure *j*, we assume that *ε*_*j*_ (measurement error for the *j*^th^ imaging variable across *i* observations) is normally distributed with zero mean and variance *V*_*j*_:

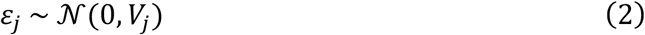

with the total residual variance 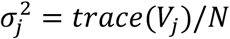. For example, given a longitudinal study design that includes family members, the random intercepts for subject and family induce a marginal variance structure *V*_*j*_ that can be parameterized as the variance attributable to living in the same family (*F*), repeated measures from the same subject (*S*), and random errors (*E*); the total normalized covariance is decomposed as:

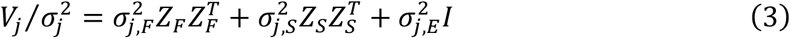

where, generically, *Z* is an indicator matrix expressing the random intercepts with row *Z*_*i*_ specifying the membership of the *i*^th^ observation and each column corresponding to a different level of the random factor, *Z*_*F*_ for families and *Z*_*S*_ for subjects, each with a corresponding variance parameter, 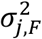 and 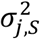, respectively; *I* is an identity matrix with dimensionality equal to number of observations and a corresponding variance parameter 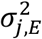, which is the variance of the random uncorrelated measurement error (or unmodeled residual variance, uncorrelated across observations). A visual example of three different experimental designs is shown in **Figure S3**.

The main computational bottleneck here is the estimation of *V*_*j*_ for every imaging outcome measure. As noted in (Gao and Owen, 2020), the cost of computing the likelihood for a given value of parameters requires *O*(*n*^3/2^) time, where *n* is the number of observations. Therefore, when the number of imaging measures, *J*, is large, such as in voxel-wise analyses, using a likelihood-based solver for every single measure would be computationally infeasible. In the following sections, we discuss how our algorithm overcomes this computational limitation.

### 2.3 Estimating the variance components

To overcome the computational bottleneck of estimating the parameterized covariance matrix, *V*_*j*_, we implement MoM estimator to obtain the values of 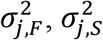, and 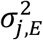 for each imaging variable *j*. The MoM estimator is fast and consistent though at the cost of statistical efficiency. Let the OLS solution for the fixed effects be:

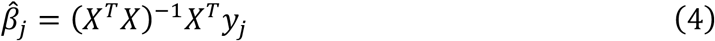

Then, the residuals can be computed as:

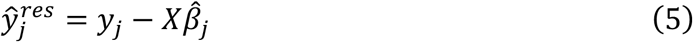

with the total residual variance 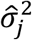 estimated as 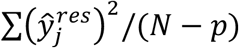. Then, for a given imaging variable *j*, and for each pair of observations (*i, i*′), the expected value of the product of corresponding residuals 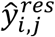 and 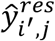 is:

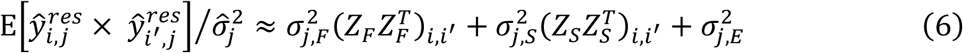

where the E operator represents the expectation with respect to random effects, and (*ZZ*^*T*^)_*i,i*_′denotes the (*i, i*′) element of the *ZZ*^*T*^ matrix (see Supplementary Materials and **Figure S1** for an intuition behind equation 6). This implies (*n* × (*n* + 1))/2 elements with three unknown parameters (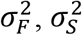, and 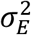), allowing us to estimate the variance parameters using a non-negativity constrained OLS (see Supplementary Materials for details on non-negativity constrained OLS and **Figure S2** for a demonstration); the values in the first column of the design matrix are determined by 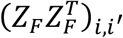, the second by 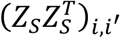, and the last column of 1’s for the error variance. Importantly, the OLS estimates can be expressed as a simple matrix expression allowing us to simultaneously obtain the variance parameters for all *J* imaging variables in a single computation. In addition to using MoM for estimating variance components, users also have the option of selecting a ML estimator, instead of the MoM estimator (see *Implementation details* in the Supplementary Materials for details).

### 2.4 Estimating fixed effects

Once the variance parameters are estimated for each *j*, the covariance matrix *V*_*j*_ can be composed as per equation (3). Then, the fixed effects can be estimated using the GLS solution:

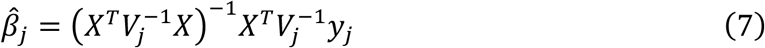

with variance

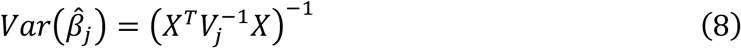

The test statistic for individual fixed effects can be computed as Wald’s ratio, 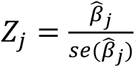 . For an arbitrary contrast estimate 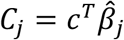, Wald’s ratio is computed using 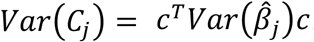. The two-tailed *p* value for the fixed effects is then calculated as:

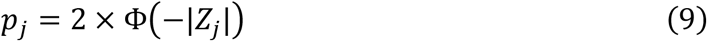

where Φ is the cumulative distribution function of a standard normal distribution, and |*Z*_*j*_| is the absolute value of the Wald’s ratio *Z*_*j*_.

To further speed up calculations, instead of separately computing *V*_*j*_ for each imaging measure, we group them on a regular multidimensional grid according to the value of estimated random effects. To clarify, depending on the grid size *K* (where *K* ≪ *J*), we create a multidimensional grid (spaced by a factor of 1/*K*). Then, using the estimated values of the variance parameters of the random effects, we bin the outcome variables – variables that have very similar random effects will have the same bin value (see *Implementation details* in the Supplementary Materials for details on binning and **Figure S4** for a visual explanation of the binning strategy). In a series of simulations, we will show that a finite number of gridded bins is sufficient to capture the variance components for all input imaging measures while dramatically reducing the required computational time.

### 2.5 Simulations

To evaluate the validity of our algorithm and demonstrate that estimates from FEMA are accurate, we performed a series of simulations. We simulated realistic data with different covariance structures for a large number of outcome variables and examined the validity and reliability of the parameter estimates and the computational time required by FEMA. We also performed comparisons with a standard mixed model implementation (*fitlme* or *fitlmematrix* in MATLAB). In all estimations involving FEMA, we used the MoM estimator for random effects and used a GLS solution for fixed effects. For all simulations, we used MATLAB R2023a (The MathWorks, Natick, USA). Note that MATLAB’s *fitlme* (and *fitlmematrix*) report the estimates and confidence intervals for random effects in standard deviation units, while FEMA reports the estimates and confidence intervals in variance units. Therefore, we squared the MATLAB derived values before comparing them with FEMA derived values. Additionally, for all variance components obtained from FEMA, we rescaled them back to their original scale before comparing them with MATLAB (i.e., the variances are reported as such and *not* as proportions of the residual variance in the data).

#### 2.5.1 Simulation 1: Binning and parameter recovery

In the first experiment, we had two objectives: first, find an optimal value of binning and second, demonstrate that the recovered parameters (slopes for the fixed effects and variance parameters of the random effects) were comparable to the parameters used for generating the data (the *ground truth*). In this simulation experiment, we generated 2000 *y* variables for 10,000 observations, with different values of family effect (*F*) and longitudinal effect (*S*) for each *y* variables. Each family was allowed to have between one and five individuals with up to five repeated observations, totaling 10,000 observations. There were five fixed effects in the simulation. The forward model was based on:

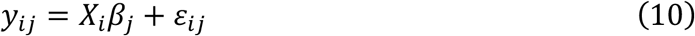

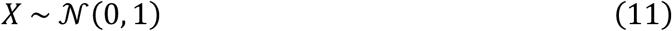

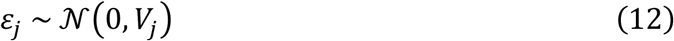

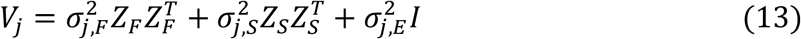

where the notations are the same as introduced in the preceding sections. To define the ground truth, we sampled the fixed effects parameters from a uniform distribution: *β*_*j*_ ∼ 𝒰[−0.02, 0.02]. The random effects were sampled from a uniform distribution, while satisfying the following conditions:

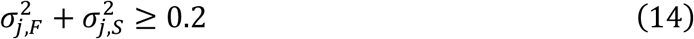

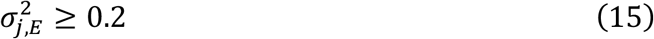

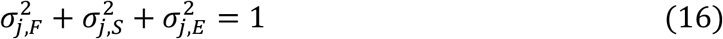

In other words, the minimum variance of the unmodeled error term (*E*) was 0.2 while the variance parameters of the other random effects ranged between 0 and 0.8. These parameter ranges were chosen based on the observations reported in large-scale imaging analyses, i.e., small effect sizes and evident effects of family and repeated measures (Dick et al., 2021). The total residual variance in the data (residual variance after removing the effect of fixed effects or the variance explained by the random effects) was set to 1.

##### 2.5.1.1 Effect of binning

Having defined the ground truth, we simulated 50 different sets of *X* and *y* variables with different underlying *Z*_*F*_ and *Z*_*S*_ structures i.e., for every simulation, a new family structure with different number of repeated observations were generated (henceforth referred as repeats and denoted with *r*). We performed parameter estimation for 20 different values of bins: five bins from 1 to 5 (spaced by 1), five bins from 10 to 50 (spaced by 10), and 10 logarithmically spaced values (rounded to their nearest integer values) between 10^2^ and 10^3.3010^, where 3.3010 is log_10_ 2000 (the number of *y* variables). For every bin, for every *y* variable, for every fixed effect *p*, we calculated the mean squared error (MSE) as the average (over the 50 repeats) of the square of the difference between the parameter estimate 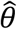 and the simulated ground truth θ^true^:

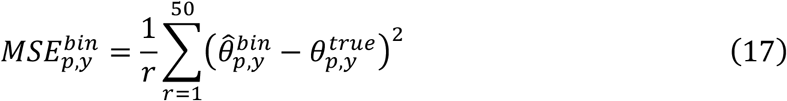

This resulted in an MSE value for each of the 20 bins, five fixed effects, and 2000 imaging variables. Then, we summed the MSE value across the 2000 imaging variables (*total MSE*) and examined this as a function of bin size. Finally, to select an optimal value for the number of bins, we averaged the total MSE (*average total MSE*) across the five fixed effects and selected a bin value where the average total MSE was comparable to the minimum of the average total MSE.

#### 2.5.1.2 Accuracy of parameter estimates

Next, we examined the accuracy of the estimated parameters from FEMA. For this comparison, we plotted (against the simulated ground truth) the estimated parameters for the five fixed effects for all 2000 imaging variables, across the 50 repeats. Additionally, we plotted the mean (over 50 repeats) estimated parameter for each fixed effect and imaging variable. Similarly, we examined the plots of estimated random effect parameters against the simulated ground truth.

### 2.5.2 Simulation 2: Comparison with *fitlme*

Next, we compared the estimated parameters from FEMA with that of a standard implementation of mixed models (*fitlme* in MATLAB; the same model can also be estimated with *fitlmematrix* function). We followed the same data generation model as simulation 1 but restricted ourselves to 50 imaging variables because of the high computational demand for solving mixed models. The simulated data for 10,000 observations included the effect of five fixed effects (an intercept term and four additional fixed effects) as well as the random effects −*F, S*, and *E*. These mixed-effects model can be specified as (note that the error term (1|*E*) is implicitly included):

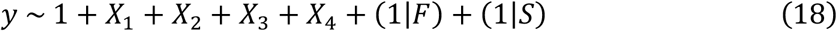

To estimate the model parameters using *fitlme*, we used the default ML estimation method. For estimating the parameters using FEMA, we set the bin value to 20. To directly compare the confidence intervals for the random effect, we additionally ran 100 permutations of wild bootstrap in FEMA and then calculated the 2.5 and 97.5 percentiles of the permuted parameter estimates to get the 95% confidence intervals on the random effects.

#### 2.5.3 Simulation 3: Performance comparison

Next, we compared the performance of FEMA with that of *fitlmematrix* in MATLAB. We opted to use *fitlmematrix* instead of *fitlme* for this set of comparison because *fitlme* requires a *table type* variable while *fitlmematrix* can work with matrices directly (similar to FEMA).

##### 2.5.3.1 Computational time as a function of number of observations

To examine the computational time required as a function of increasing number of observations, we varied the sample size from 2000 to 20,000 observations (in steps of 2000 observations) keeping the number of imaging variables fixed at 50. The number of fixed effects were five. As before, we only used one bin value of 20 when running FEMA but did not perform any permutations to estimate the confidence intervals on the random effects. We followed a similar data generation scheme as simulation 1 with the key difference that the data generation model included the genetic relatedness matrix (GRM) as an additional random effect. To clarify, some of the variance in the data was now additionally attributed to how (genetically) similar individuals were within a family. In this synthetic data situation, the genetic relationship within the family was centered at 0.5 with a small amount of noise and the genetic relationship between repeated observations would be 1. The rationale for including the additive genetic variance, *A* effect as a random effect was so that we could showcase the computational time needed by FEMA under a variety of common model specifications of varying levels of complexity. Having already established the accuracy of the model parameters, for this analysis we were primarily interested in comparing the computational time necessary for estimating different model parameters. Therefore, we generated the data from a *F, A, S*, and *E* specification and then recovered the parameters (using *fitlmematrix* in MATLAB and FEMA) for: a) *F* and *E* ; b) *S* and *E* ; and c) *F, S*, and *E* random effect specification. Additionally, we recovered the parameters using FEMA for the following random effects specifications: a) *A* and *E*; b) *F, A*, and *E*; c) *S, A*, and *E*; and d) *F, A, S*, and *E*. For the latter four models, we did not specify the models in MATLAB because it is uncertain whether a continuous valued *N* × *N* matrix can be specified as a single random effect using *fitlmematrix*^1^.

We note that several functions implemented in MATLAB are multithreaded by default. The benefits of multithreaded operation will apply to the functions internally called by MATLAB’s *fitlmematrix*, as well as FEMA. However, one could argue that conventional mixed models could be sped up further by using parallel processing. Therefore, in addition to comparing the time required by FEMA and *fitlmematrix* (when serially looping over *y* variables), we also calculated the time needed for *fitlmematrix* using parallel computing. When using parallel processing, one needs to ensure that there is an optimal usage of resources so that there is no bottleneck. To ensure a (reasonably) fair comparison between FEMA and *fitlmematrix* (with and without parallel processing), we ran the entire simulation as a *slurm* cluster job on a high-performance cluster (HPC). To clarify, the calls to FEMA and *fitlmematrix* (with and without parallel processing) were part of the same script that was run on the HPC with the following settings: CPUs per task = 40 and memory per CPU = 20 GB (the HPC cluster consisted of AMD EPYC™ 7702 64-core processors, resulting in 128 available cores with a maximum memory availability of 3955 GB, out of which we used a maximum of 40 CPUs and 20 GB of RAM per CPU). Further, when initializing a parallel pool in MATLAB, we set the number of parallel workers to 20. When using parallel processing in MATLAB, MATLAB defaults to 1 thread per worker. We changed this specification to 2 threads per worker for the timing comparison. In addition, since parallel processing would invoke multiple MATLAB instances in the background (which would remain idle when using FEMA or doing serial processing), we closed the parallel pool immediately after the parallel processing was completed (for each number of observation). The reported time taken does not include the timing overhead for creating or closing a parallel pool, overheads associated with parsing the returned MATLAB model to extract relevant parameters, or time taken for initializing or saving variables.

##### 2.5.3.2 Computational time as a function of number of imaging variables

We additionally examined how the computational time increased as a function of increasing number of imaging variables. For this simulation, we fixed the number of observations to 10,000. The number of fixed effects were five and the random effects were *F, S*, and *E*. We varied the number of imaging variables from 100 to 5000 (step size of 100 variables). Similar to the previous comparison, we calculated the time taken for estimation of the parameters for fixed and random effects by FEMA (bin value of 20), *fitlmematrix*, and a call to *fitlmematrix* wrapped in a parallel *for* loop (with the same cluster settings as before).

#### 2.5.4 Simulation 4: Type I error rate

Having demonstrated parameter recovery (via simulation 1), general similarity to a standard mixed model solver (via simulation 2), and computational efficiency (via simulation 3), we examined the type I error rate for fixed effects using FEMA (bin value of 20) and compared it with MATLAB’s *fitlmematrix*. Specifically, we simulated 10,000 total observations with 100 fixed effects and *F, S*, and *E* random effects for 50 imaging variables. The variance parameters for the random effects were the same as simulation 1 but the slopes for the fixed effects were set to zero (i.e., the simulated data had no contribution from the *X* variables). Then, we estimated the slopes for the fixed effects and the variance components for the random effects using FEMA and *fitlmematrix*. We repeated this process 100 times. For every repeat, we counted the number of times the *p* values for the fixed effects were smaller than four threshold values of *α =* 0.05, 0.01, 0.001, and 0.0001, for every outcome variable. Then, for every repeat, we summed the number of false positives (FPs) across the outcome variables to get total number of FPs for that repeat (at that threshold value of *α*). Finally, to get an aggregate number across the 100 repeats, we calculated the mean of the total number of false positives and rounded it towards positive infinity (i.e., a *ceil* function). Since there were 50 imaging variables with 100 fixed effects each, at each of these threshold values, we did not expect (on average) more than 250, 50, 5, and 1 FPs (for *α =* 0.0001, the number of FPs is 1 because we are rounding the average towards positive infinity).

#### 2.5.5 Simulation 5: Effect of number of observations

As an additional analysis, we examined the effect of number of observations on parameter recovery for both the fixed effects and the random effects. Since FEMA is designed for large sample size situations, we examined how the estimates from FEMA converged to the estimates from an ML-based solver. Specifically, for this simulation, we examined parameter recovery as a function of 16 increasing sample sizes: 50, 10 sizes between 100 and 1,000 (spaced by 100), and five sizes between 2,000 and 10,000 (spaced by 2000). Note that here we are interchangeably using the terms *sample size* and *number of observations*. Similar to other simulations, we specified five fixed effects with true effect uniformly distributed between -0.2 and 0.2; there were two random effects – family effect and subject effect, and the simulated data had up to five family members within a family and up to five repeated observations for a given subject. We simulated 500 outcome variables and ran five repeats of this experiment (i.e., for every repeat, for every sample size, we created fresh draws of fixed and random effects). For each of these simulated datasets, we fit LME models using FEMA (at a bin value of 20) and ML-estimator using *fitlmematrix* function in MATLAB. Then, for every sample size, for every repeat, we examined the squared difference between the estimates from FEMA and the ground truth; similarly, we estimated the squared differences between the estimates from *fitlmematrix* and the ground truth. Then, we averaged these squared differences across the five repeats and summed these squared differences across the 500 outcome variables. We similarly performed the same calculations for the variance parameter estimates for the random effects.

#### 2.5.6 Simulation 6: Type I error rate as a function of bin value

As a final simulation, we examined the rate of FPs as a function of the bin value. We simulated 10,000 observations with 100 fixed effects and *F, S*, and *E* random effects for 500 imaging variables. The variance parameters for the *F* and *S* random effects were set to be between 0.3 and 0.4 – this narrow range ensured that several outcome variables would be evaluated together (i.e., they would be *binned* together for computing *V* when GLS solution for fixed effects is implemented), thereby allowing us to examine if binning had any adverse effect on FPs. Similar to simulation 4, when simulating the data, we set the slopes for the fixed effects to zero (i.e., the simulated data had no contribution from the *X* variables). Then, we estimated the slopes for the fixed effects using FEMA at bin values 1 to 30. We repeated this process 100 times. For every repeat, for every bin value, we counted the number of times the *p* values for the fixed effects were smaller than *α =* 0.05 for every outcome variable. Then, for every repeat, for every bin value, we summed the number of FPs across the outcome variables to get total number of FPs for that repeat and that bin size. Finally, we averaged this number (rounded to positive infinity) across repeats to get an average number of FPs for each bin value. Since there were 500 imaging variables with 100 fixed effects each, we did not expect (on average) more than 2500 FPs at *α =* 0.05.

### 2.6 Empirical application

To demonstrate the utility of FEMA, we applied FEMA to the ABCD Study^®^ (data release 4.0) to examine the longitudinal changes in imaging measures during adolescence. The ABCD Study^®^ is a longitudinal study of brain development in adolescents in the United States. The ABCD Study^®^ protocols were approved by the University of California, San Diego Institutional Review Board and data was collected after taking informed consent from the parent/caregiver’s as well as child’s assent. The multimodal imaging acquisition in the ABCD Study^®^ includes T1- and T2-weighted structural scans, multi-shell diffusion scans, and resting and task fMRI (see, (Casey et al., 2018) for details on image acquisition and (Hagler et al., 2019) for details on image processing and analyses methods). Release 4.0 includes two scans –one at baseline and the second scan after two years from the baseline.

Most standard neuroimaging analyses can be formulated to be one of the following categories –voxel-wise analyses (e.g., NIfTI images), vertex-wise analyses (e.g., GIfTI images), ROI-wise analyses (tabular data), or connectome-wide analyses (high dimensional flattened tabular data). In addition, most non-imaging variables can be structured as tabular data type with rows being observations and columns being the variable(s) of interest. To show the use-cases of FEMA for these data types, we present the results from the longitudinal analyses of two measures – cortical thickness derived from T1-weighted structural images (ROI-wise and vertex-wise analyses) and connectome-wide functional connectivity (high dimensional flattened tabular data). We have previously demonstrated the applicability of FEMA to voxel-wise data, analyzing restricted spectrum imaging parameters derived from diffusion weighted images (Palmer et al., 2022), and to non-imaging tabular data, analyzing cognitive variables (Smith et al., 2023). These analyses were run on MATLAB R2023a on a server having 48 Intel® Xeon® E5-2680 CPUs, 12 CPUs per socket (2 sockets), and 2 threads per core; the server had 512 GB of RAM. Prior to running the analyses, we standardized each *y* variable to have zero mean and unit standard deviation. Similar to the simulation experiments, we squared the *fitlmematrix* derived standard deviation values before comparing them with FEMA derived variance values. Additionally, we report all variances from FEMA as such and *not* as proportions of the residual variance in the data).

#### 2.6.1 Outline of image processing

As mentioned previously, details of the ABCD multimodal image acquisition can be found in (Casey et al., 2018) and a detailed description of image processing pipeline can be found in (Hagler et al., 2019). Additionally, the current analyses use data from ABCD data release 4.0; the details of changes between the image processing pipeline as reported in (Hagler et al., 2019) for data release 2.0.1 and data release 4.0 can be found in modality specific documentation with the release notes at the National Institute of Mental Health Data Archive (NDA): https://nda.nih.gov/study.html?id=1299. For completeness, we provide a brief overview of the structural and functional preprocessing below.

Broadly, the T1-weighted (T1w) images were corrected for gradient distortion using scanner-specific non-linear transformations, followed by bias correction using a novel bias correction algorithm. The bias corrected T1w was then segmented using Freesurfer 7.1.1 and different morphometric features like cortical thickness computed for different atlases. For the purpose of this study, we used the average cortical thickness across 68 ROIs from the Desikan-Killiany atlas (Desikan et al., 2006).

For resting state fMRI data, the first step in the preprocessing pipeline was to perform motion correction using AFNI’s (Cox, 1996) *3dvolreg*. Next, the motion corrected images were corrected for B0 distortion using FSL’s TOPUP v5.0.2.2 (Andersson et al., 2003; Smith et al., 2004), followed by gradient distortion correction (Jovicich et al., 2006). The distortion corrected images were then rigidly aligned with T1w images based on registration of the field maps with T1w images. Following these, dummy scans and first few time points were removed, and voxel-wise time series were normalized by the voxel-specific mean of the time series (i.e., each voxel’s time series was scaled by that voxels’ average BOLD signal over time). The normalized time series was demeaned and multiplied by 100. The denoising step consisted of performing a linear regression on the time series with the independent variables being the quadratic trends, six motion parameters (their derivatives and squares), mean time series from white matter, ventricles, and the whole brain, and their first order derivates. Additionally, the motion time courses were temporally filtered to attenuate signals linked to respiration (Fair et al., 2020). Time points which had a framewise displacement > 0.3 mm were excluded during regression (Power et al., 2014). These values were replaced by linear interpolation (after regression and residualization), followed by band-pass filtered between 0.009 – 0.08 Hz (Hallquist et al., 2013). This denoised time series was then projected to subject-specific cortical surfaces and ROI-specific time series was calculated as the average (across vertices) time series within each ROI. For the purpose of this study, we used the Fisher transformed (inverse hyperbolic tangent) functional connectivity matrices within and between the Desikan-Killiany (Desikan et al., 2006), the Destrieux (Destrieux et al., 2010), and Gordon (Gordon et al., 2016) cortical parcellation schemes, as well as selected subcortical regions from the Freesurfer “aseg” whole brain segmentation (Fischl et al., 2002) (see section Connectome-wide functional connectivity).

#### 2.6.2 Cortical thickness

##### 2.6.2.1 ROI-level cortical thickness

For the first demonstration using real data, we used the average cortical thickness across 68 ROIs from the Desikan-Killiany atlas (Desikan et al., 2006) and compared the estimates obtained from FEMA (at a bin value of 20) with those obtained from using *fitlmematrix* in MATLAB. The goal of this analysis was to show that the parameter estimates at a bin size of 20 were similar to the ones obtained from a standard implementation. We calculated the confidence intervals for random effects using 100 permutations of wild bootstrap. For this analysis, we only considered those subjects who had an image at both the baseline as well as the follow-up and that both images passed quality assurance (i.e., the variable *imgincl_t1w_include*, defined in the NDA data dictionary, was equal to 1). The sample size was 6314 subjects (in each visit) across 5387 families. The cross-sectional age was between 107 months and 133 months (mean ± SD = 118.82 ± 7.45 months) and the follow-up age was between 127 months and 166 months (mean ± SD = 143.28 ± 7.78 months).

The fixed effects (covariates) included: intercept, the age of the participants at the time of recruitment (*age*_*recruitment*_, in months), the change in age between baseline and the follow-up visit (*age*_*delta*_), dummy-coded sex variable (two levels; retained one level), dummy-coded scanner device number (31 levels; retained 29 levels), dummy-coded scanner software version number (17 levels; retained 15 levels), dummy-coded household income level (three levels; retained two levels), dummy-coded parental education level (five levels; retained four levels), and the first 20 genetic principal components (which would account for the population structure in the data) derived using the GENESIS package (Gogarten et al., 2019). In total, there were 74 covariates out of which the covariates of interest were *age*_*recruitment*_ and *age*_*delta*_. Note that we dropped two levels from the various dummy-coded variables so that the matrix of covariates was not rank deficient. In addition, to model the within-family and within-subject variances (not explained by the fixed effects), we specified the *F* and *S* random effects. This model (for each imaging variable) can be written as (note that the error term (1|*E*) is implicitly included):

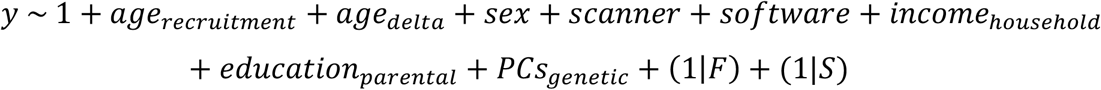

where, *age*_*recruitment*_ would remain the same for baseline visit as well as follow-up visit, while *age*_*delta*_ would be zero for baseline visit and be equal to the difference between the *age*^*followUp*^ and *age*^*recruitment*^ .

##### 2.6.2.2 Vertex-wise cortical thickness

Using the same model as for the ROI-level analyses, we examined the vertex-wise association of cortical thickness with *age*^*recruitment*^ and *age*_*delta*_. The sample size was the same as the ROI-level analysis and the total number of analyzable vertices were 18,742.

#### 2.6.3 Connectome-wide functional connectivity

For the second demonstration of the use of FEMA, we analyzed the resting state functional connectivity between 582 ROIs, resulting in a total of 169,071 pairs of connectivity values (see (Hagler et al., 2019) for details on the denoising of the time series and subsequent calculation of the connectivity values). These 582 ROIs consisted of 68 ROIs from the Desikan-Killiany parcellation scheme (Desikan et al., 2006), 148 ROIs from the Destrieux parcellation scheme (Destrieux et al., 2010), 333 ROIs from the Gordon parcellation scheme (Gordon et al., 2016), and 33 ROIs from the aseg parcellation scheme (Fischl et al., 2002) (not including lesion and vessal parcellations). The Gordon parcellation scheme further consisted of 13 network communities – default, somatomotor hand and mouth, visual, frontoparietal, auditory, cingulo-parietal and cingulo-opercular, retrosplenial temporal, ventral and dorsal attention, salience, and none. For performing connectome-wide analysis, we followed the same model as cortical thickness. We only included subjects who had a resting state scan at baseline and at the follow-up visit and that both the images passed quality assurance (i.e., the variable *imgincl_rsfmri_include*, defined in the NDA data dictionary, was equal to 1). The sample size was 4994 subjects (in each visit) across 4375 families and the number of connectivity values were 169,071. The cross-sectional age was between 107 months and 133 months (mean ± SD = 119.10 ± 7.52 months) and the follow-up age was between 127 months and 166 months (mean ± SD = 143.58 ± 7.86 months).

## 3 Results

### 3.1 Simulation 1: Binning and parameter recovery

#### 3.1.1 Effect of binning

For every bin value, we calculated the total MSE across 2000 imaging variables for the five fixed effects. Then, we calculated the average MSE across these five fixed effects to get an overall estimate of the total MSE for each bin value. We observed that the minimum average of the total MSE was at a bin value of 100 (total MSE = 0.13247); however, the MSE at a bin value of 20 was comparable to this minimum (total MSE = 0.13255), at a fraction of the computational time required (**Figure 1**). Therefore, we opted to report all subsequent results at a bin value of 20.

**Figure 1:**
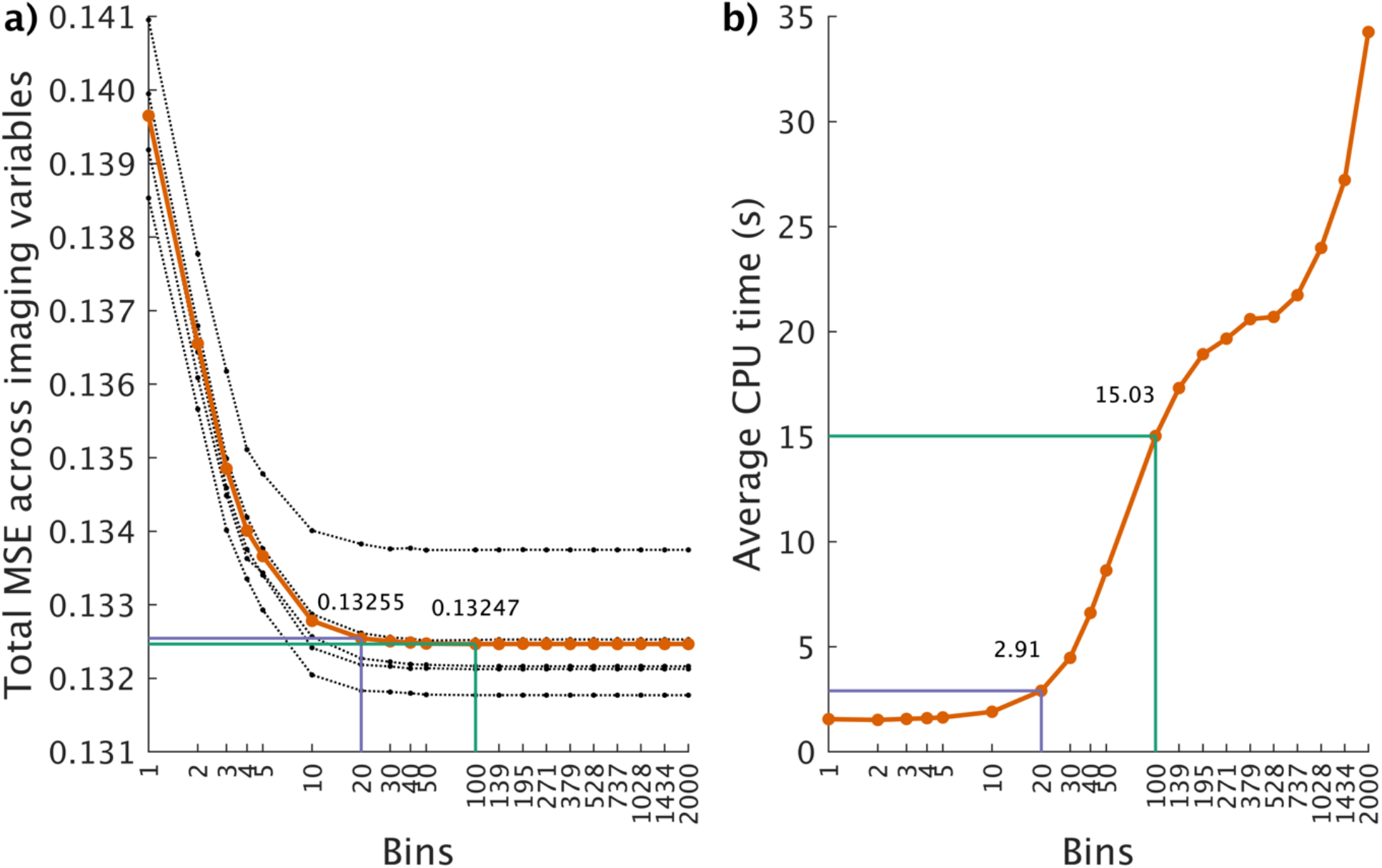
The effect of binning on mean squared error (MSE) of parameter estimates of fixed effects. We simulated 2000 imaging variables with five fixed effects (10,000 observations). Then, using 20 different bin values, we estimated the parameters using FEMA and calculated the mean (over 50 repeats) squared error of the parameter estimates. The dotted black lines in panel **a)** indicate the total MSE (across the 2000 imaging variables) for each of the five fixed effects while the solid orange line indicates the average of this total MSE across the five fixed effects. We observed that the minimum total MSE was at a bin value of 100 (indicated by the green line) while a bin value of 20 (indicated by the purple line) showed a comparable MSE; panel **b)** shows the computational time required for each bin value (averaged across 50 repeats); the computational time for bin value of 20 (purple line) is a fraction of computational time required for bin value of 100 (green line). Note that the *x*-axis is non-linear for both the panels.

#### 3.1.2 Accuracy of fixed effects and random effects

For each of the fixed effects, the average (over 50 repeats) estimated parameters were comparable to the ground truth (see **Figure 2a**). The five fixed effects showed a similar pattern of variation around the mean with no obvious outliers. Similarly, for the random effects, we observed comparable average (over 50 repeats) estimated parameters to the simulated ground truth (see **Figure 2b**). Between the three random effects, the family effect *F* had the least amount of variation in the estimated parameters while the subject effect *S* and the unmodeled error term *E* had relatively larger variation in the estimated parameters across the 50 repetitions. We did note that the family effect *F* had relatively larger variation around the ground truth at larger values.

**Figure 2:**
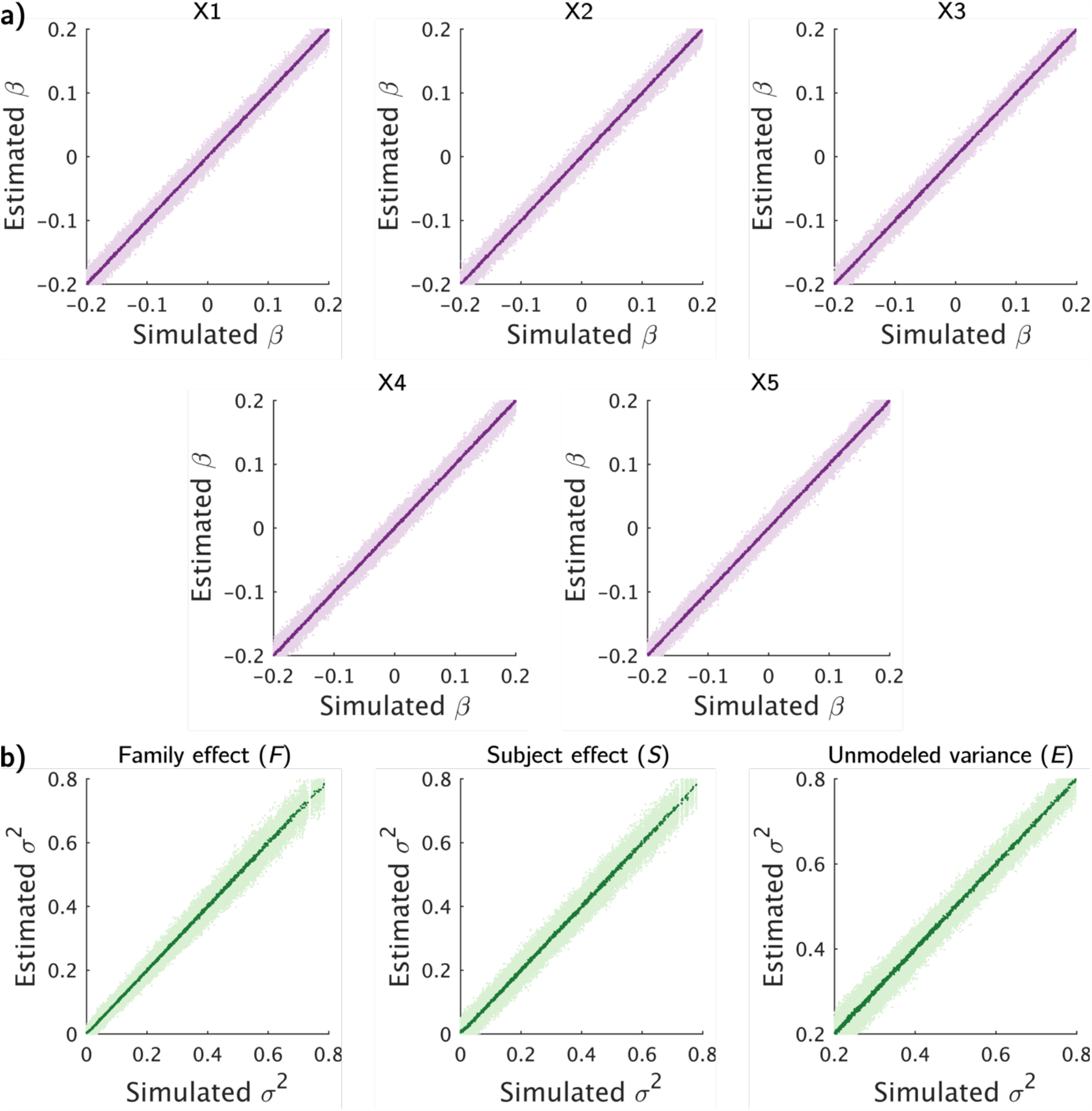
Comparison of estimated parameters against the simulated ground truth for: **a)** fixed effects, and **b)** for random effects. The light purple and light green dots indicate the estimated parameters for 2000 imaging variables at 50 different repetitions for the fixed and random effects respectively, while the dark purple and dark green dots indicate the average (over 50 repetitions) of these parameters for the 2000 imaging variables for the fixed and random effects respectively.

### 3.2 Simulation 2: Comparison with *fitlme*

When we compared the fixed effects estimates from FEMA to that obtained from calling the *fitlme* function in MATLAB, we found that the results were identical with similar point estimates and standard errors for all imaging variables for all five fixed effects (**Figure S5**). When examining the random effects, the point estimates for all imaging variables for the three random effects were comparable between FEMA and *fitlme*, albeit some differences in the 95% confidence interval range, which is likely because of different methods of estimation of the confidence intervals (non-parametric wild bootstrap in FEMA vs. ML-based calculation in *fitlme*; **Figure S6**). We additionally quantified the squared difference between the parameter estimates from FEMA and *fitlme* and summed it across fixed (or random) effects and outcome imaging variables. This total squared difference for the fixed effects was 9.2573e-06 showing that the point estimates were identical, and for the random effects was 7.14 showing small differences in the point estimates (as is visually evident from examining **Figure S6**).

### 3.3 Simulation 3: Performance comparison

#### 3.3.1 Computational time as a function of number of observations

We compared the time taken by FEMA and MATLAB to estimate model parameters (slopes for the fixed effects and variance components of the random effects) for 50 imaging variables as a function of number of observations. Generally speaking, FEMA was several times faster than the *fitlmematrix* implementation in MATLAB. For the *F* and *E* model specification, FEMA was between 3.8 to 12.6 times faster than MATLAB’s *fitlmematrix* and between 2.1 to 8.7 times faster than the *fitlmematrix* called in parallel (**Figure 3a**). For the *S* and *E* model, FEMA was between 5.6 to 9.5 times faster than *fitlmematrix* and between 1.3 and 2.4 times faster than *fitlmematrix* called in parallel (**Figure 3b**), and for the *F, S*, and *E* model, FEMA was between 14 to 27.1 times faster than *fitlmematrix* and between 2.2 and 3.7 times faster than *fitlmematrix* called in parallel (**Figure 3c**). Therefore, in all simulation circumstances, FEMA outperformed a standard LME solver in terms of computational time, and the computational speed of FEMA (run without parallel processing) was evident even when the standard LME solver was called in a parallel computing environment. When we examined the time taken by FEMA for other random effects configuration, we found a similar trend with the maximum time of about 5.3 seconds for 20,000 observations (for the *F, A, S*, and *E* model) (**Figure S7**). Therefore, even after including the additive genetic effect as a random effect in FEMA, we did not see a substantial increase in computational time for a large number of imaging variables.

**Figure 3:**
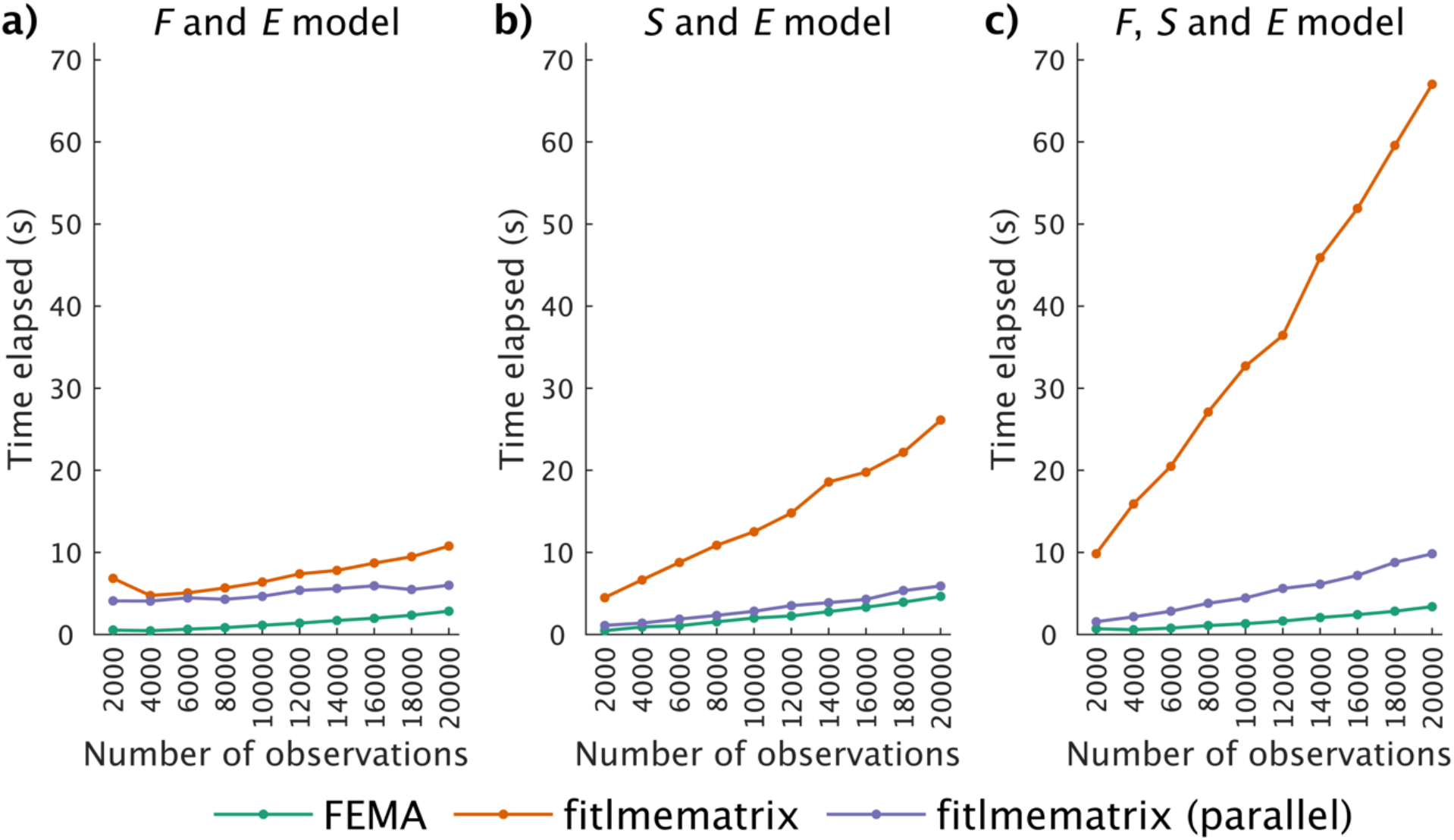
Comparison of time taken to fit different models using FEMA (green), *fitlmematrix* in MATLAB (orange), and implementation of *fitlmematrix* using parallel computing (purple) as a function of increasing number of observations for 50 imaging variables and five fixed effects. The random effects included in each model are indicated at the top of each panel: family effect *F*, subject effect *S*, and the unmodeled variance term *E*.

#### 3.3.2 Computational time as a function of number of imaging variables

When we examined the computational time needed by FEMA and MATLAB as a function of increasing number of imaging variables, we observed that the amount of time taken by FEMA was more or less steady while the time taken by MATLAB increased linearly with increasing number of imaging variables (**Figure 4**). This is likely due to the fact that *fitlmematrix* is not designed to handle multiple imaging variables and therefore there is a computational overhead when repeatedly calling *fitlmematrix* using a loop; this is not the case with FEMA because it is designed to handle a large number of outcome variables with vectorized operations that scale well to a large number of outcome variables. We also found a similar linear increase in computational time when calling *fitlmematrix* in parallel (**Figure 4**), with FEMA being several times more efficient (between 40.2 and 1020.6 times faster than *fitlmematrix* and 7.3 and 125 times faster than *fitlmematrix* being called in parallel). Extrapolating from this linear model, the time taken by *fitlmematrix* to fit a mixed-effects model for 558,718 voxels for 10,000 observations would be about 5.4 days, while parallel implementation of *fitlmematrix* would take about 15.4 hours. On the other hand, the extrapolated timing for FEMA to fit the same model would be about 3.4 minutes.

**Figure 4:**
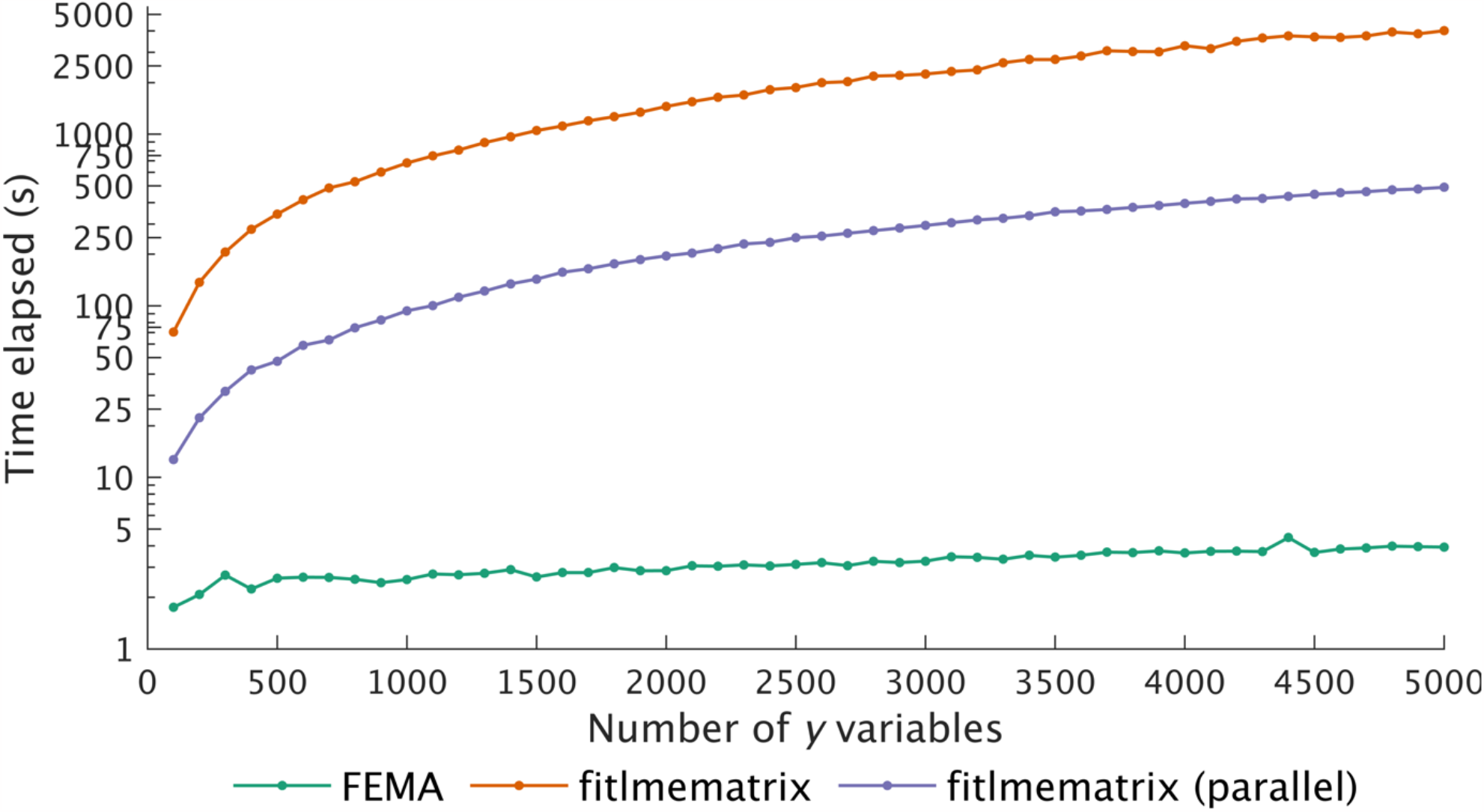
Comparison of time taken by FEMA (green), MATLAB’s *fitlmematrix* (orange), and implementation of *fitlmematrix* using parallel processing (purple) as a function of increasing number of imaging variables for fitting a model with 10,000 observations, five fixed effects, and the family effect *F*, subject effect *S*, and the unmodeled variance term *E*, specified as random effects. Note that the *y*-axis is non-linear.

### 3.4 Simulation 4: Type I error rate

On comparing the type I error rate for fixed effects estimation between a standard mixed model solver and FEMA, we observed comparable number of FPs across different threshold values of false positive rates, demonstrating that FEMA did not have any inflation of FPs. We observed comparable values to the expected average number of FPs (see **Table 1**) and these numbers were similar to the ones reported by *fitlmematrix* (see **Figure S8** for a detailed look at the number of FPs at 5% false positive rate threshold).

**Table 1:** Comparison of type I error rate for fixed effects estimation between FEMA and *fitlmematrix* function from MATLAB; we performed 100 repetitions of simulating 10,000 observations for 50 imaging variables, with each imaging variable associated with 100 *X* variables (having zero effect); family and subject effects were specified as random effects. At each threshold of false positive (FP), we counted the number of times we observed a statistically significant *p* value at that threshold and summed it across the 50 imaging variables to get total FP for every repeat; the reported counts are averages of the total FPs across 100 repeats, rounded to nearest infinity (i.e., using *ceil* function).

| | Avg. #FP $\alpha = 0.05$ | Avg. #FP $\alpha = 0.01$ | Avg. #FP $\alpha = 0.001$ | Avg. #FP $\alpha = 0.0001$ |
| --- | --- | --- | --- | --- |
| **FEMA** | 246 | 49 | 5 | 1 |
| *fitlmematrix* | 258 | 53 | 6 | 1 |

### 3.5 Simulation 5: Effect of number of observations

When examining the differences between ground truth and estimated parameters from FEMA and *fitlmematrix* as a function of number of observations, we found that the fixed effects estimates converged to ML-based estimates at a few hundred observations (**Figure S9**), while for the random effects variance parameter estimates, FEMA estimates converged to ML-based estimates at a few thousand observations (**Figure S10**).

### 3.6 Simulation 6: Type I error rate as a function of bin values

On examining the type I error rate, we did not find any influence of the bin value on the number of FPs. On average, for every bin, the number of FPs was comparable and did not exceed the 5% FP rate threshold (**Figure S11**).

### 3.7 Empirical application

#### 3.7.1 Cortical thickness

##### 3.7.1.1 ROI-level cortical thickness

When examining the ROI-level cortical thickness data, we found comparable slope estimates and comparable 95% confidence intervals between MATLAB’s *fitlmematrix* and FEMA (at a bin value of 20) for both the cross-sectional effect of age (*age*^*recruitment*^, **Figure S12**) as well as the longitudinal effect of in age (*age*_*delta*_, **Figure S13**). Although the random effects were not of interest in this study, we compared the variance estimates and the confidence intervals of the random effects with *fitlmematrix* derived estimates for completeness. The *p*-values for most ROIs survived a Bonferroni correction for multiple comparison correction (*α =* 0.05/68) except for bilateral temporal pole, left superior temporal, bilateral precuneus, right parahippocampal, bilateral entorhinal, and left caudal middle frontal (for *age*_*recruitment*_), and bilateral entorhinal and right temporal pole (for *age*_*delta*_). Similarly, for random effects, the FEMA estimated variance components and their confidence interval values were comparable to the estimates from MATLAB (**Figure S14**). The general pattern showed (relatively) small variances being explained by the family effect and a larger contribution (albeit with larger confidence intervals) of the subject effect for all ROIs. The unmodeled variance term was relatively small for most regions.

##### 3.7.1.2 Vertex-wise cortical thickness

Vertex-wise analysis of cortical thickness revealed widespread negative effect of both cross-sectional effect of age as well as longitudinal effect of age on cortical thickness. The peak negative effect of age was in the bilateral rostral middle frontal regions and the left isthmus cingulate region. A small fraction of vertices (812 vertices out of the total 18,742) showed a small positive effect of age and these were primarily located in the bilateral precentral regions (**Figure S15**). The longitudinal effect of age was prominently distributed across most of the cerebral cortex and was predominantly negative, with the largest negative effect size in bilateral cuneus and precuneus regions. A small number of the vertices (349 vertices out of the total 18,742) showed a positive effect of the longitudinal effect of age and these were primarly located in the bilateral precentral regions (**Figure 5**). Performing whole-brain vertex-wise analysis (for 6314 subjects with two data points and across 18,742 vertices) with *F, S*, and *E* model random effects specification with FEMA took about 11 seconds.

**Figure 5:**
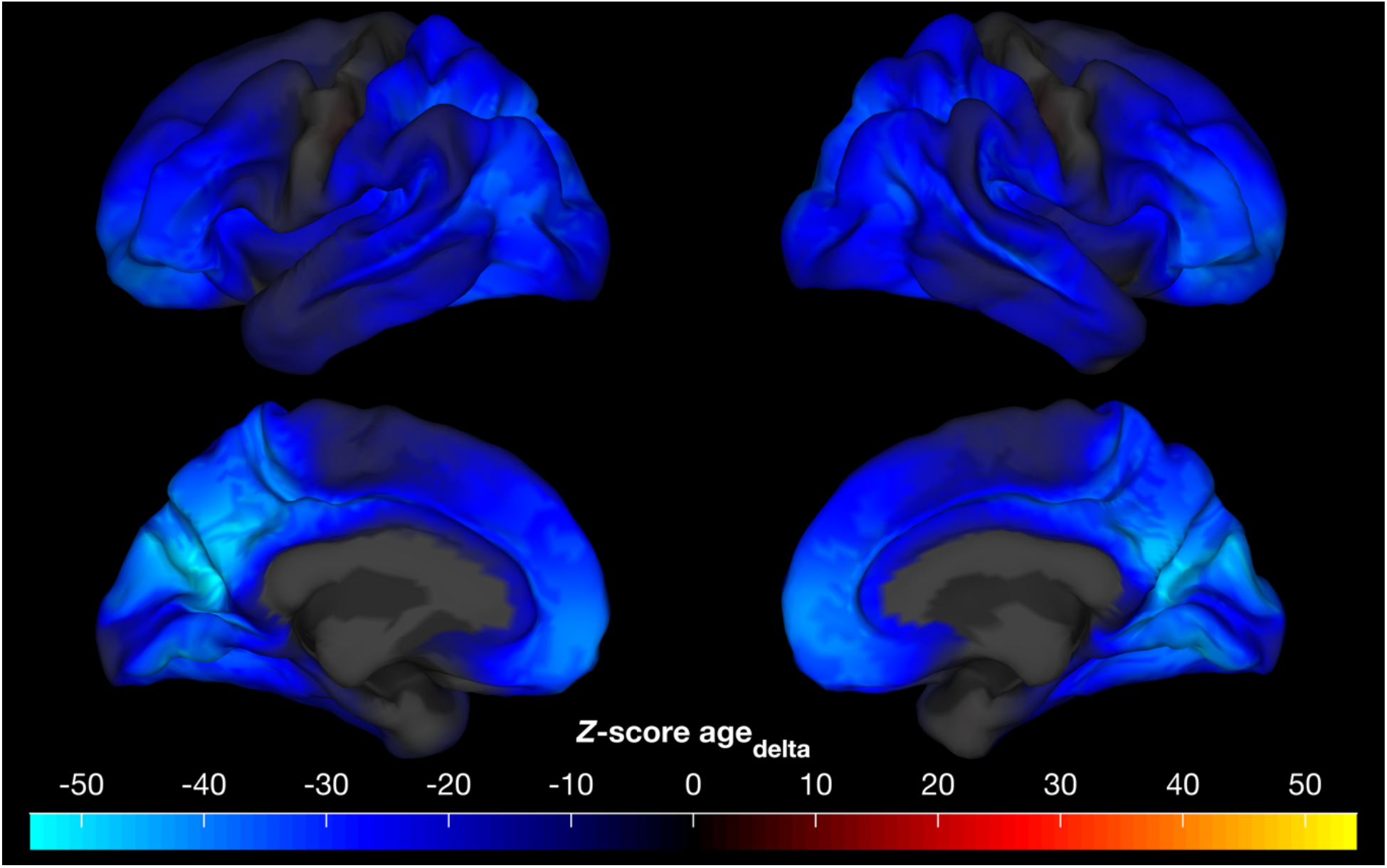
Unthresholded vertex-wise *Z* scores for the longitudinal effect of age (*age*_*delta*_), performing whole-brain vertex-wise analysis using FEMA took about 11 seconds.

#### 3.7.2 Connectome-wide functional connectivity

On examining the effect of age on connectome-wide pairs of connectivity values derived from resting state fMRI, we found both positive and negative effects of the cross-sectional effect of age, although the effects were small with the *Z* scores ranging between -4.05 and 4.78. About 55% of the 169,071 pairs of connectivity values showed a positive effect of the cross-sectional effect of age and the effects were well distributed across the atlases and parcellation communities (**Figure S16**). On the other hand, the longitudinal effect of age was more pronounced in both the positive and the negative direction, with *Z* scores ranging between - 23.70 and 20.07. About 59% of the 169,071 pairs of connectivity values showed a negative effect of the longitudinal effect of age with effects well distributed within and across the atlases/parcellation communities (**Figure 6**). Performing connectome-wide analysis (for 4994 subjects with two data points and across 169,071 pairs of connectivity values) with *F, S*, and *E* random effects specification with FEMA took about 54 seconds.

**Figure 6:**
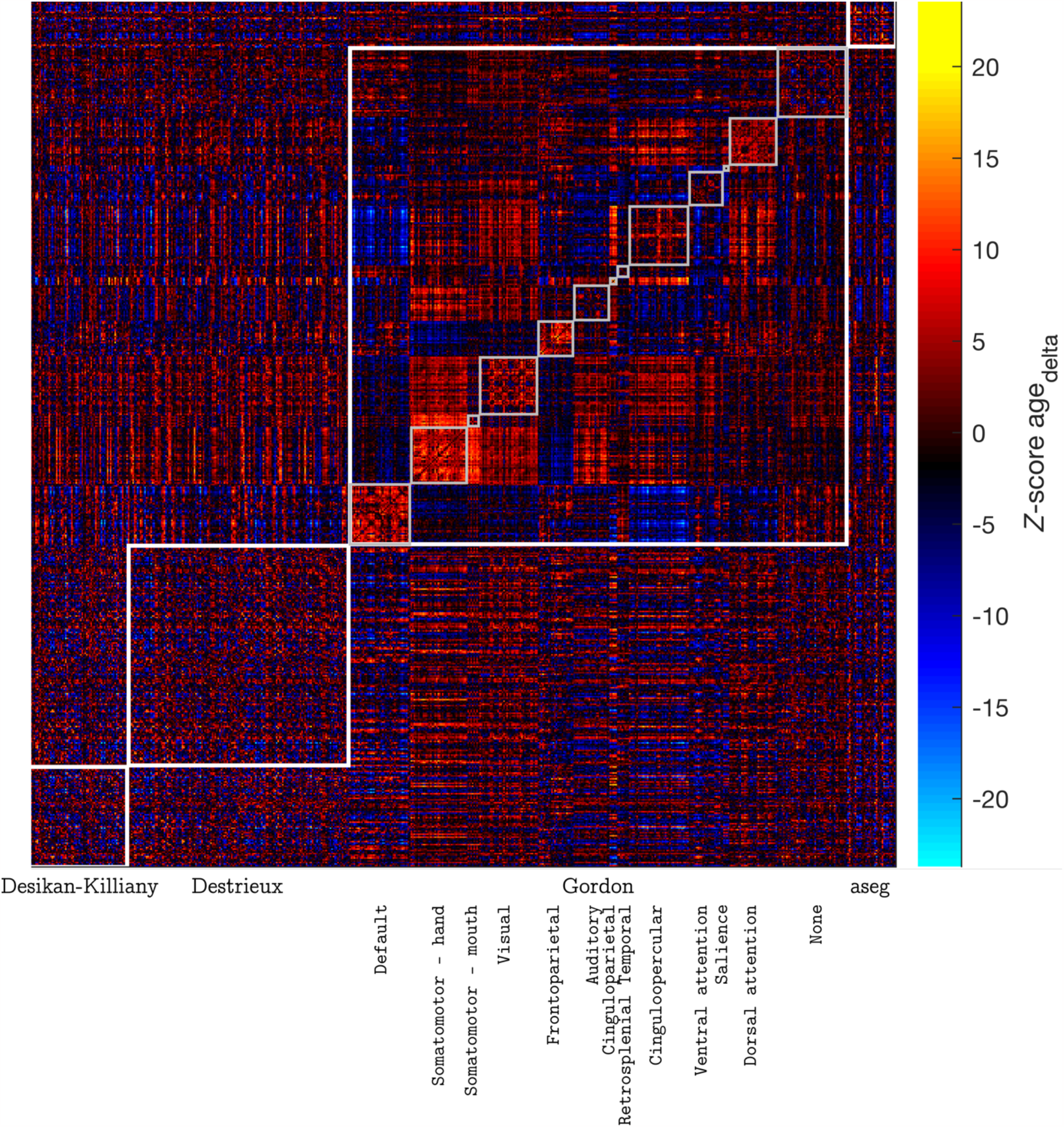
Distribution of *Z*-scores for the longitudinal effect of age (*age*_*delta*_) across 582 ROIs; the ROIs were defined using a combination of Desikan-Killiany atlas, Destrieux atlas, Gordon parcellation, and the aseg atlas; the Gordon parcellation is further divided into 13 communities; performing connectome-wide analysis using FEMA took about 54 seconds.

## 4 Discussion

In this paper, we have introduced a novel method, FEMA, for performing mixed-effects analyses for whole-brain large-scale samples. In the first part of the paper, we have described the problem statement and how FEMA finds the solution to estimating model parameters in a computationally efficient manner. Next, through extensive simulations, we have shown that the estimates (for both fixed effects as well as random effects) obtained from FEMA are comparable to simulated ground truth as well as estimates obtained from a standard ML-based solver, at a fraction of computational time. Importantly, these simulations revealed that the computational performance of FEMA was efficient even when compared against a standard LME solver called within a parallel computing environment. Further, we have shown that FEMA performs these estimations without any inflated type I error rate. Finally, in the third part of this paper, we have demonstrated three applications of FEMA – first, at an ROI level, where we examined the cross-sectional and longitudinal effect of age on cortical thickness; second, at whole-brain vertex-wise level, of the effect of age on cortical thickness; and finally, at connectome-wide level, of the effect of age on functional connectivity derived from resting state functional MRI. These demonstrate the applicability of FEMA on three types of data formats – summary tabulated data, surface data (or GIfTI images), and high-dimensional tabular data (connectome-wide). In addition, as previously mentioned, we have demonstrated the use of FEMA with voxel-wise data or NIfTI images (Palmer et al., 2022), and non-imaging tabular data (Smith et al., 2023).

FEMA achieves computational efficiency owing to the following factors: i) implementation of a sparse random effects design; ii) an efficient binning strategy that speeds up estimation of fixed effects; iii) using a method of moments estimator for random effects; and iv) use of vectorized operations for scalability across variables. While this is not the first time that method of moments has been used for estimating model parameters in a mixed model setting (see, for example, (Gao, 2017; Gao and Owen, 2017, 2020) and some of the references cited therein), to the best of our knowledge, this is the first time that such a scalable solution has been presented in the context of solving mixed models for neuroimaging data in large samples. Our solution additionally circumvents the need for parallel computing or high memory requirements. Through the results of simulation 1, we have shown that using a small number of bins (such as bin value of 20) is adequate for the accurate estimation of the fixed and random effects, while keeping the computational time low. Further, through simulations 1 and 2, we have shown that these estimates are similar to the simulated ground truth as well as comparable to estimates from a standard ML-based mixed model solver. In terms of computational efficiency, through simulation 3, we have shown that the time required by FEMA is a fraction of the time taken by MATLAB’s built-in functions. Even when engaging parallel computing for MATLAB’s *fitlmematrix*, the time taken by FEMA was always lesser than MATLAB, demonstrating the benefits of the implementation in FEMA. In addition, the results of simulation 3 also shows that the time taken by FEMA does not grow appreciably with increasing number of imaging variables (a benefit of vectorized operations that scale well to multiple imaging variables), thereby allowing fast and efficient whole brain analyses to be completed in a matter of seconds to minutes as opposed to prohibitively long computational time using a standard solver. We also note that FEMA is computationally efficient even when adding genetic relatedness as a random effect. While it may appear (from **Figure 3** and **Figure 4**) that it may be possible to exploit parallel processing to an extent that solving whole brain mixed models no longer takes a long time, we highlight two aspects of this comparison: one that when benchmarking the performances, we purposefully used a highly powered parallel configuration with a large amount of computing resources with 20 parallel workers and 2 threads per worker and large memory resources; this is more computational power than is available to many groups and thus depending on the computational resources that a researcher has (and other factors like whether there are additional processes running), this time may be substantially lower than practically realizable for many labs. Second, running a large number of parallel jobs is also memory intensive and thereby requires the availability of such computing facility. Recently, (Maullin-Sapey and Nichols, 2022) developed the “Big” Linear Mixed Models (BLMM) toolbox which uses a combination of novel closed form solutions, vectorized computing, and distributed processing on clusters (while keeping the memory requirement in check) for solving mixed models. While we have not compared the computational performance of FEMA with BLMM, we highlight that FEMA is fast and computationally efficient and one could complete whole brain analyses on a regular laptop/workstation within a short time (as long as the data fits in the memory), thereby circumventing having dedicated computing clusters for such analyses. Finally, we note that these computational advantages do not come at the cost of an inflated type I error, as demonstrated by the results of simulation 4 and simulation 6, where we saw that the average type I error rate was controlled at 5%.

Applying FEMA to real ABCD Study^®^ data revealed interesting patterns of the cross-sectional and longitudinal effect of age on cortical thickness and functional connectivity values. The comparative results from our application of FEMA to the ROI-level cortical thickness data and fitting the same model in MATLAB confirmed (in conjunction with the simulation results) that the parameter estimates from FEMA (at a bin value of 20) were accurate and similar to the estimates using an ML estimator in MATLAB. On further examining the vertex-wise distribution of the effect of age, we found widespread negative effect of both the cross-sectional effect of age as well as longitudinal effect of age. Similar findings of the negative effect of age on cortical thickness has been previously reported (see, for example, (Brown and Jernigan, 2012) for a review). A recent comprehensive analysis of cortical thickness changes across the lifespan found that cortical thickness across most brain regions achieved a maximum value between 3 and 10 years of age, followed by a decline in cortical thickness (Frangou et al., 2022). However, these finding is not without contradiction and as reviewed and discussed in (Walhovd et al., 2017), multiple factors can lead to differing results. We hope that FEMA applied to large-scale dataset will allow careful modeling of various fixed and random effects, leading to more consistent findings in a dramatically shorter time. In contrast to cortical thickness, when we applied FEMA to connectome-wide pairs of functional connectivity values, we found both positive and negative effect of cross-sectional and longitudinal effect of age. A recent study (Edde et al., 2021) synthesized the functional connectivity literature across the human lifespan and found that adolescence was marked with more subtle changes in brain organization. It was also interesting to note that both cortical thickness and functional connectivity values had a stronger association with longitudinal age than with cross-sectional age. Since the goal of the current manuscript was merely to demonstrate the utility of FEMA, we have not provided an in-depth examination of these fixed effects or examined the spatial patterns of different random effects. Examining and comparing these patterns of changes in the brain during these transitory adolescent periods will lead to insights into the neurobiology of brain development and may possibly also shed new light on normal and pathological brain trajectories.

### 4.1 Limitations and future directions

FEMA allows analyzing mixed models in a computationally efficient manner that opens new possibilities of performing large-scale studies at fine-grained voxel-level or vertex-level data. The FEMA code is highly parallelizable where computation across bins can be spread across multiple parallel workers, thereby allowing almost real-time processing (see *Implementation details* section in the supplementary materials). We also note that FEMA has wrapper functions allowing for permutation testing using the PALM toolbox (Winkler et al., 2014) and calculation of TFCE statistics (Smith and Nichols, 2009) for both voxel-wise and vertex-wise data. However, our work is not without its limitations. First, FEMA uses a moment-based estimator and therefore is based on the law of large numbers. While we have performed a simulation examining the effect of number of observations on parameter recovery and our simulations suggest that a few hundred samples are adequate for having comparable fixed effect estimates to ML-based solver (**Figure S9**), we cannot give a general recommendation on how large sample sizes should be, as this would depend on the intricacies and nuances of the data itself. Since our implementation of FEMA additionally includes ML-based estimation, it can be considered as a safer choice, albeit at the cost of increased computational time. Second, the current implementation of FEMA assumes that the outcome variables are continuous variables (i.e., FEMA currently cannot, for example, account for categorical outcome variables). Third, in its current form, we do not account for the covariances between different random effects (although it is possible to do so). Fourth, we only estimate the variance components of the random effects and not the estimates of best linear unbiased predictors of random effects at each level of the grouping variables. Finally, we note that the calculation of the *p* values for the fixed effects is based on the standard normal distribution – we assume that since the sample size is large, there would be no difference between the *p* values derived from a standard normal distribution vs. a *T* distribution with *n* − *k* degrees of freedom (where *n* is the sample size and *k* are the number of fixed effects; this ‘residual’ degrees of freedom is the default method of calculating *p* values in MATLAB). However, this issue is not without differing points of view and we point the readers to (Kuznetsova et al., 2017; Maullin-Sapey and Nichols, 2022, 2021) and the references cited therein for a discussion on degrees of freedom and calculating *p* values in linear mixed models.

Future work will explore the application of FEMA on large scale datasets like the ABCD Study^®^ and UK Biobank where we will explore different modeling strategies and hope to uncover new insights about the changes in multimodal properties of the brain. We have previously demonstrated the equivalence of FEMA and OpenMX and applied FEMA on cognitive variables from the ABCD Study^®^ (Smith et al., 2023). Similarly, in (Palmer et al., 2022), we have shown the applicability of FEMA on voxel-wise age-related changes in white matter and subcortical regions. In a different use-case of FEMA, in (Zhao et al., 2023), we show how the beta estimates for the fixed effects from FEMA could be used for the next steps in the study (in this case, for eventual prediction of behavioral outcome). Depending on the study dataset, future FEMA applications will additionally incorporate genetic similarity between the study participants, account for the shared in-utero environment in twins, and other factors like shared home environment; these, and examination of the spatial patterns of different random effects are exciting avenues to explore, that will form the basis of future FEMA applications. In addition, we are currently incorporating another estimator for random effects into FEMA. This is the iterative GLS (IGLS) estimator proposed by Goldstein (Goldstein, 1986). The formulation is similar to equation (4) with the key difference that the random effects are estimated using the GLS solution (which we use for estimating the fixed effects). The procedure is repeated (hence the name iterative) until the coefficients do not change appreciably. Goldstein (Goldstein, 1986) showed that in the normal case, using IGLS would be equivalent to using ML for estimation of the random effects. This method is slower than using a single iteration of MoM but is faster than using ML. Further details on the IGLS can be found in various sources (see, for example, (Goldstein, 2004, chap. 2, 1986)) and details of a REML equivalent of the IGLS, the restricted IGLS, can be found in (Goldstein, 1989). We also note, in passing, that an implementation of the IGLS was previously used in the context of neuroimaging variables in (Lindquist et al., 2012). We are also working on extending FEMA to perform genome-wide association analyses, which will enable performing streamlined imaging-genetics analyses. In parallel, FEMA is being packaged and deployed as part of the Data Exploration and Analysis Portal (DEAP) 2.0, which will enable users to examine and perform analyses on the ABCD Study^®^ data via a graphical user interface. Additionally, deploying the parallel version of FEMA on DEAP 2.0 will speed up the computation to the extent that it will enable users to test hypotheses in real time. Finally, FEMA is available to the public as a MATLAB-based toolbox with added applications that allow volume and surface visualizations – combining this with the Multi-Modal Processing Stream (MMPS) package (Hagler et al., 2019) will allow the users to have an end-to-end pipeline that allows image preprocessing, mixed model analyses, as well as visualization of results.

### 4.2 Usage notes

Here, we briefly recap the most salient information for using FEMA (more detailed description of these points can be found under the heading *Implementation details* in the Supplementary Materials). Currently, the following random effects are supported within FEMA: family effect, subject effect, additive genetic effect, dominant genetic effect, maternal effect, paternal effect, twin effect, and the home effect. Within FEMA, there are two estimators available for estimating these variance components: a MoM estimator and a ML estimator. Generally speaking, if the random effects by themselves are not particularly of interest, we suggest using MoM estimator as it will be significantly faster to compute, without sacrificing the accuracy of the fixed effects estimates; on the other hand, if the random effects are of interest, a user may want to use ML to estimate the variance parameters of the random effects. Additionally, the accuracy of fixed effects will be determined by the number of bins. As shown in the results on simulated data (**Figure S5**) and from the results of ROI-level cortical thickness data (**Figures S12** and **S13**), a bin value of 20 is a reasonable choice that balances computational time and accuracy of the fixed effects. On a related note, the number of iterations can, theoretically speaking, improve the estimation accuracy of the coefficients, although in practice we have only seen minor changes in the estimates with higher number of iterations. A final usage information is the reporting of the random effects – we report the variance components as proportions of the total residual variance and separately report the total residual variance. Therefore, interested users have the option of scaling the variance components to their original estimated values.

## 5 Data/code availability

The source code for FEMA is available to the public via GitHub: https://github.com/cmig-research-group/cmig_tools/; the ABCD Study^®^ data is available via the NIMH Data Archive: https://nda.nih.gov/abcd/. The codes used for analyses and creating the figures in this paper are available at: https://github.com/parekhpravesh/FEMA.

## 6 Acknowledgements

This work was supported by grant R01MH122688, RF1MH120025, and R01MH118281 funded by the National Institute for Mental Health (NIMH). Data used in the preparation of this article were obtained from the Adolescent Brain Cognitive Development^SM^ (ABCD) Study (https://abcdstudy.org), held in the NIMH Data Archive (NDA). The ABCD Study^®^ is supported by the National Institutes of Health and additional federal partners under award numbers U01DA041048, U01DA050989, U01DA051016, U01DA041022, U01DA051018, U01DA051037, U01DA050987, U01DA041174, U01DA041106, U01DA041117, U01DA041028, U01DA041134, U01DA050988, U01DA051039, U01DA041156, U01DA041025, U01DA041120, U01DA051038, U01DA041148, U01DA041093, U01DA041089, U24DA041123, U24DA041147. A full list of supporters is available at https://abcdstudy.org/about/federal-partners/. A listing of participating sites and a complete listing of the study investigators can be found at https://abcdstudy.org/wp-content/uploads/2019/04/Consortium_Members.pdf. ABCD consortium investigators designed and implemented the study and/or provided data but did not necessarily participate in the analysis or writing of this report. This manuscript reflects the views of the authors and may not reflect the opinions or views of the NIH or ABCD consortium investigators. The ABCD data repository grows and changes over time. The ABCD data used in this report came from http://dx.doi.org/10.15154/1523041. The fast track data release used in this report are available at https://nda.nih.gov/edit_collection.html?id=2573. Instructions on how to create an NDA study are available at https://nda.nih.gov/nda/tutorials/creating_an_nda_study).

PP acknowledges funding from the European Union’s Horizon 2020 research and innovation programme under the Marie Skłodowska-Curie grant agreement number 801133 and from the Research Council of Norway grant number 324252.

Parts of this work used the TSD (Tjeneste for Sensitive Data) facilities, owned by the University of Oslo, operated and developed by the TSD service group at the University of Oslo, IT-Department (USIT), using resources provided by UNINETT Sigma2 –the National Infrastructure for High Performance Computing and Data Storage in Norway.

Some of the color schemes used for various figures were taken from ColorBrewer (Brewer, n.d.). Some of the figures use the tight_subplot function (Pekka Kumpulainen (2023). tight_subplot(Nh, Nw, gap, marg_h, marg_w) (https://www.mathworks.com/matlabcentral/fileexchange/27991-tight_subplot-nh-nw-gap-marg_h-marg_w), MATLAB Central File Exchange. Retrieved April 5, 2023).

## 7 Conflict of interest

Dr. Anders M. Dale was a founder of and holds equity in CorTechs Labs, Inc, and serves on its Scientific Advisory Board. He is also a member of the Scientific Advisory Board of Human Longevity, Inc. (HLI), and the Mohn Medical Imaging and Visualization Centre in Bergen, Norway. He receives funding through a research agreement with General Electric Healthcare (GEHC). The terms of these arrangements have been reviewed and approved by the University of California, San Diego in accordance with its conflict-of-interest policies. Dr. Andreassen has received speaker fees from Lundbeck, Janssen, Otsuka, and Sunovion and is a consultant to Cortechs.ai. The other authors declare no competing interests.

## 8 Author contributions (CRediT roles)

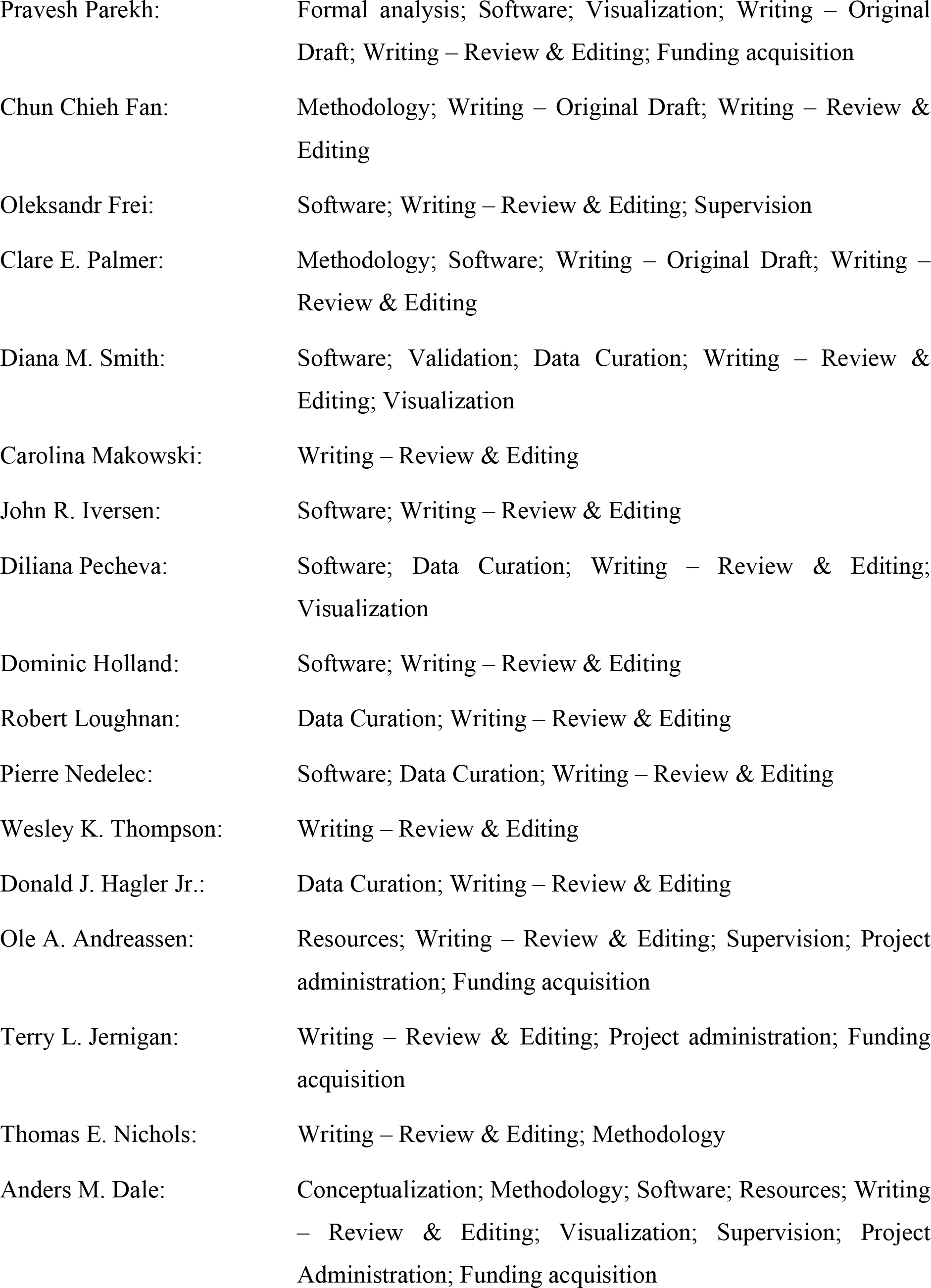

8. Author contributions (CRediT roles):
|  |  |
| --- | --- |
| Pravesh Parekh: | Formal analysis; Software; Visualization; Writing – Original Draft; Writing – Review & Editing; Funding acquisition |
| Chun Chieh Fan: | Methodology; Writing – Original Draft; Writing – Review & Editing |
| Oleksandr Frei: | Software; Writing – Review & Editing; Supervision |
| Clare E. Palmer: | Methodology; Software; Writing – Original Draft; Writing – Review & Editing |
| Diana M. Smith: | Software; Validation; Data Curation; Writing – Review & Editing; Visualization |
| Carolina Makowski: | Writing – Review & Editing |
| John R. Iversen: | Software; Writing – Review & Editing |
| Diliana Pecheva: | Software; Data Curation; Writing – Review & Editing; Visualization |
| Dominic Holland: | Software; Writing – Review & Editing |
| Robert Loughnan: | Data Curation; Writing – Review & Editing |
| Pierre Nedelec: | Software; Data Curation; Writing – Review & Editing |
| Wesley K. Thompson: | Writing – Review & Editing |
| Donald J. Hagler Jr.: | Data Curation; Writing – Review & Editing |
| Ole A. Andreassen: | Resources; Writing – Review & Editing; Supervision; Project administration; Funding acquisition |
| Terry L. Jernigan: | Writing – Review & Editing; Project administration; Funding acquisition |
| Thomas E. Nichols: | Writing – Review & Editing; Methodology |
| Anders M. Dale: | Conceptualization; Methodology; Software; Resources; Writing – Review & Editing; Visualization; Supervision; Project Administration; Funding acquisition |

## Footnotes

1. We note that it may be possible to specify such a matrix using the *fitlmematrix* function in MATLAB. However, preliminary testing in this regard was deemed unsuccessful, as the computation never finished even after several days. We also note that it may be possible to reparametrize the GRM which might make the computation tractable, although we have not examined this aspect (see, for example, (Hunter, 2021; McArdle and Prescott, 2005; Rabe-Hesketh et al., 2008; Wang et al., 2011))

## Notes

### Summary of Updates

Timing updates; additional simulation

