## Supplementary Materials for "FEMA: Fast and efficient mixed-effects algorithm for large sample whole-brain imaging data"

**List of authors:**

Pravesh **Parekh**[1,*,#], Chun Chieh **Fan**[2,3,#], Oleksandr **Frei**[1,4], Clare E. **Palmer**[5], Diana M. **Smith**[5,6,7,8], Carolina **Makowski**[3,6], John R. **Iversen**[5,9,10], Diliana **Pecheva**[3,6], Dominic **Holland**[3], Robert **Loughnan**[11], Pierre **Nedelec**[12], Wesley K. **Thompson**[2], Donald J. **Hagler** Jr.[3,6], Ole A. **Andreassen**[1], Terry L. **Jernigan**[3,5,13,14], Thomas E. **Nichols**[15,16], Anders M. **Dale**[3,6,13,14,17,*]

**Affiliations:**

[1]NORMENT, Division of Mental Health and Addiction, Oslo University Hospital & Institute of Clinical Medicine, University of Oslo, Oslo, Norway

[2]Center for Population Neuroscience and Genetics, Laureate Institute for Brain Research, Tulsa, OK, USA

[3]Department of Radiology, School of Medicine, University of California San Diego, La Jolla, CA, USA

[4]Centre for Bioinformatics, Department of Informatics, University of Oslo, Oslo, Norway

[5]Center for Human Development, University of California San Diego, La Jolla, CA, USA

[6]Center for Multimodal Imaging and Genetics, University of California San Diego, La Jolla, CA, USA

[7]Neurosciences Graduate Program, University of California San Diego, La Jolla, CA, USA

[8]Medical Scientist Training Program, University of California San Diego, La Jolla, CA, USA

[9]Institute for Neural Computation, University of California San Diego, La Jolla, CA, USA

[10]The Swartz Center for Computational Neuroscience, University of California San Diego, La Jolla, CA, USA

[11]Population Neuroscience and Genetics Lab, University of California San Diego, La Jolla, CA, USA

[12]Department of Radiology and Biomedical Imaging, University of California San Francisco, San Francisco, CA, USA

[13]Department of Cognitive Science, University of California San Diego, La Jolla, CA, USA

[14]Department of Psychiatry, University of California San Diego, La Jolla, CA, USA

[15]Big Data Institute, Li Ka Shing Centre for Health Information and Discovery, Nuffield Department of Population Health, University of Oxford, Oxford, UK

[16]Wellcome Centre for Integrative Neuroimaging, FMRIB, Nuffield Department of Clinical Neurosciences, University of Oxford, Oxford, UK

[17]Department of Neuroscience, University of California San Diego, La Jolla, CA, USA

[#] Equal contribution

**Correspondence should be addressed to:*

**Pravesh Parekh**,
NORMENT, Oslo University Hospital,
Ullevål Hospital, building 48,
Kirkeveien 166,
Oslo 0450, Norway


**Prof. Anders M. Dale**,
University of California,
San Diego School of Medicine,
9500 Gilman Drive,
La Jolla, CA 92037, USA,


### 1. Materials and Methods

#### 1.1 Estimating the variance components

As mentioned in the main text, let the ordinary least squares (OLS) solution for the fixed effects be:

$$\hat{\beta}_j = (X^T X)^{-1} X^T y_j \tag{1}$$

Then, the residuals can be computed as:

$$\hat{y}_j^{res} = y_j - X\hat{\beta}_j \tag{2}$$

with the total residual variance $\hat{\sigma}_j^2$estimated as $\sum\left(\hat{y}_j^{res}\right)^2/(N-p)$. Then, for a given voxel $j$, and for each pair of observations $(i, i')$, the expected value of the product of corresponding residuals $\hat{y}_{i,j}^{res}$ and $\hat{y}_{i',j}^{res}$ is:

$$\mathrm{E}\left[\hat{y}_{i,j}^{res} \times \hat{y}_{i',j}^{res}\right]/\hat{\sigma}_j^2 \approx \sigma_{j,F}^2 (Z_F Z_F^T)_{i,i'} + \sigma_{j,S}^2 (Z_S Z_S^T)_{i,i'} + \sigma_{j,E}^2 \tag{3}$$

where the E operator represents the expectation with respect to random effects, and $(ZZ^T)_{i,i'}$ denotes the $(i, i')$ element of the $ZZ^T$ matrix.

To gain an intuition about this equation, consider a case of a single voxel. The left side of the equation, the product of $\hat{y}_i^{res}, \hat{y}_{i'}^{res}$, is the squares (when $i = i'$) and products (when $i \neq i'$, but only for instances where $i$ and $i'$ have a known source of correlation; see section *Sparsity of the random effects* in the *Implementation details*) of the residuals, written out as a vector. The right side of the equation are the squares and the products of the elements of the $ZZ'$ matrix, written out in vector forms for $(i, i')$ entries. This implies $(n \times (n+1))/2$ elements with three unknown parameters ($\sigma_F^2$, $\sigma_S^2$, and $\sigma_E^2$), allowing us to estimate the variance parameters using a non-negativity constrained OLS; the values in the first column of the design matrix are determined by $(Z_F Z_F^T)_{i,i'}$, the second by $(Z_S Z_S^T)_{i,i'}$, and the last column of 1's for the error variance ($\sigma_F^2$, $\sigma_S^2$, and $\sigma_E^2$). We demonstrate this using a toy example in **Figure S1**.

The non-negative least squares estimation is performed using a modified version of the Lawson-Hanson algorithm (Lawson and Hanson, 1995). While the algorithm is generic, we note that in the present context, the predictor variables are the columns constructed from the squares and products of the elements of $ZZ'$ matrix (one column for each random effect), the outcome variable is the squares and products of the $\hat{y}_i^{res}, \hat{y}_{i'}^{res}$ term, and the parameters to be estimated are the variance components for each of the random effect. To estimate these parameters with a non-negativity constrain, we first perform a least squares estimation and find the outcome variables which have at least one negative estimated parameter (variance component). Then, for every combination of random effects, we re-estimate the least squares solution by iteratively dropping random effects. To clarify, consider each random effect to have two states (zero and non-zero). Then, there would be $2^z$ configurations of these random effects (where, $z$ is the total number of random effects). If the state is zero, then that particular random effect is eliminated from the model and the model parameters re-estimated using OLS, with an additional constrain that we take the maximum value between zero and the estimated value (i.e., negative estimates are set to zero). This gives us estimates for the variance parameters over all $2^z$ configurations ranging from the scenario of considering all random effects to eliminating all random effects from the model. For each of these configurations, we calculate the sum of squared differences between the actual value of the outcome variable, and its predicted value (the predicted value is calculated as the product of columns of design matrix and the estimated non-negative variance parameters). The random effect configuration where this sum of squared differences ("cost") is minimum, is then selected as the solution. We have provided a demonstration of this algorithm using a toy example in **Figure S2**.

| Entry | Family | Subject | $y^{res}$ |
|---|---|---|---|
| $y_{F01S01r1}$ | F01 | S01 | −30 |
| $y_{F01S01r2}$ | F01 | S01 | −25 |
| $y_{F01S02r1}$ | F01 | S02 | −20 |
| $y_{F01S02r2}$ | F01 | S02 | −15 |
| $y_{F01S03r1}$ | F01 | S03 | −25 |
| $y_{F01S03r2}$ | F01 | S03 | −20 |
| $y_{F01S04r1}$ | F01 | S04 | −15 |
| $y_{F01S04r2}$ | F01 | S04 | −10 |
| $y_{F02S05r1}$ | F02 | S05 | 10 |
| $y_{F02S05r2}$ | F02 | S05 | 20 |
| $y_{F02S06r1}$ | F02 | S06 | 15 |
| $y_{F02S06r2}$ | F02 | S06 | 25 |
| $y_{F02S07r1}$ | F02 | S07 | 20 |
| $y_{F02S07r2}$ | F02 | S07 | 30 |
| $y_{F02S08r1}$ | F02 | S08 | 15 |
| $y_{F02S08r2}$ | F02 | S08 | 25 |

| Products of entries | $y_i^{res} \times y_{i'}^{res}$ | F | S | E | Term |
|---|---|---|---|---|---|
| $y_{F01S01r1} \times y_{F01S01r1}$ | −30 × −30 = 900 | 1 | 1 | 1 | $\sigma_F^2 + \sigma_S^2 + \sigma_E^2$ |
| $y_{F01S01r1} \times y_{F01S01r2}$ | −30 × −25 = 750 | 1 | 1 | 0 | $\sigma_F^2 + \sigma_S^2$ |
| $y_{F01S01r1} \times y_{F01S02r1}$ | −30 × −20 = 600 | 1 | 0 | 0 | $\sigma_F^2$ |
| $y_{F01S01r1} \times y_{F01S02r2}$ | −30 × −15 = 450 | 1 | 0 | 0 | $\sigma_F^2$ |
| $y_{F01S01r1} \times y_{F01S03r1}$ | −30 × −25 = 750 | 1 | 0 | 0 | $\sigma_F^2$ |
| ⋮ | ⋮ | ⋮ | ⋮ | ⋮ | ⋮ |
| $y_{F01S01r2} \times y_{F01S01r2}$ | −25 × −25 = 625 | 1 | 1 | 1 | $\sigma_F^2 + \sigma_S^2 + \sigma_E^2$ |
| $y_{F01S01r2} \times y_{F01S02r1}$ | −25 × −20 = 500 | 1 | 0 | 0 | $\sigma_F^2$ |
| ⋮ | ⋮ | ⋮ | ⋮ | ⋮ | ⋮ |
| $y_{F01S04r2} \times y_{F01S04r2}$ | −10 × −10 = 100 | 1 | 1 | 1 | $\sigma_F^2 + \sigma_S^2 + \sigma_E^2$ |
| $y_{F02S05r1} \times y_{F02S05r1}$ | 10 × 10 = 100 | 1 | 1 | 1 | $\sigma_F^2 + \sigma_S^2 + \sigma_E^2$ |
| $y_{F02S05r1} \times y_{F02S05r2}$ | 10 × 20 = 200 | 1 | 1 | 0 | $\sigma_F^2 + \sigma_S^2$ |
| $y_{F02S05r1} \times y_{F02S06r1}$ | 10 × 15 = 150 | 1 | 0 | 0 | $\sigma_F^2$ |
| ⋮ | ⋮ | ⋮ | ⋮ | ⋮ | ⋮ |
| $y_{F02S08r1} \times y_{F02S08r2}$ | 15 × 25 = 375 | 1 | 1 | 0 | $\sigma_F^2 + \sigma_S^2$ |
| $y_{F02S08r2} \times y_{F02S08r2}$ | 25 × 25 = 625 | 1 | 1 | 1 | $\sigma_F^2 + \sigma_S^2 + \sigma_E^2$ |

$\sigma_F^2 + \sigma_S^2 + \sigma_E^2$ = $\mathrm{E}[\hat{y}_i^{res}, \hat{y}_{i'}^{res}]$ when $F$ = 1, $S$ = 1 & $E$ = 1 = 437.50

*and* $\sigma_F^2 + \sigma_S^2$ = $\mathrm{E}[\hat{y}_i^{res}, \hat{y}_{i'}^{res}]$ when $F$ = 1, $S$ = 1 & $E$ = 0 = 406.25

*So,* $\sigma_F^2$ = $\mathrm{E}[\hat{y}_i^{res}, \hat{y}_{i'}^{res}]$ when $F$ = 1, $S$ = 0 & $E$ = 0 = 392.71

$\therefore \sigma_S^2$ = 406.25 – 392.71 = 13.54

*and* $\sigma_E^2$ = 437.50 – 406.25 = 31.25

**Figure S1:** Implementing the method of moments-based estimation of the variance parameters of the random effects. Consider an example where there are observations collected from two families $F01$ and $F02$. Within each family, there are four subjects with two observations $r_1$ and $r_2$; $S01$ to $S04$ in $F01$ and $S05$ to $S08$ in $F02$. Let the residuals (obtained after estimating the fixed effect coefficients using ordinary least squares – equations 1 and 2) be as denoted in column $y^{res}$. Then, equation 3 can be implemented as denoted in the column "Product of entries". This translates into squares and products of $y^{res}$. Note that these calculations are performed within each family, and no between family term is included. Similar to writing out these squares and products of residuals, the indicator variables $Z_F$ and $Z_S$ can be written (indicated here as columns $F$ and $S$). Since the unmodeled variance or the error term $E$ is independently distributed, the $E$ column has a value of 1 only when squares of residuals are considered (i.e., when the same entry is multiplied with itself). These squares and products of residuals can be understood as sums of different variance components, indicated in the "Term" column. The lower part of the figure shows the calculation of the variance parameters for the random terms, once they have been written out as squares and products of residuals.

**a)** Let the squares and products of residuals and design matrix be:

| $y_i^{res} \times y_{i'}^{res}$ | *F* | *S* | *E* |
|---|---|---|---|
| 25 | 1 | 1 | 1 |
| 50 | 1 | 1 | 0 |
| 75 | 1 | 1 | 0 |
| 100 | 1 | 1 | 1 |
| 150 | 1 | 1 | 0 |
| 225 | 1 | 1 | 1 |
| 400 | 1 | 1 | 1 |
| 500 | 1 | 0 | 0 |
| 625 | 1 | 1 | 1 |

Then, the unconstrained ordinary least squares solution for variance parameters can be calculated as:

| | | | | |
|---|---|---|---|---|
| $\sigma_F^2 + \sigma_S^2 + \sigma_E^2$ | = | $\mathrm{E}[\hat{y}_i^{res}, \hat{y}_{i'}^{res}]$ when $F$ = 1, $S$ = 1 & $E$ = 1 | = | 275.00 |
| and $\sigma_F^2 + \sigma_S^2$ | = | $\mathrm{E}[\hat{y}_i^{res}, \hat{y}_{i'}^{res}]$ when $F$ = 1, $S$ = 1 & $E$ = 0 | = | 91.67 |
| So, $\sigma_F^2$ | = | $\mathrm{E}[\hat{y}_i^{res}, \hat{y}_{i'}^{res}]$ when $F$ = 1, $S$ = 0 & $E$ = 0 | = | 500.00 |
| $\therefore \sigma_S^2$ | = | 91.67 − 500.00 | = | −408.33 |
| and $\sigma_E^2$ | = | 275.00 − 91.67 | = | 183.33 |

Estimated variance parameters:

$\sigma_F^2$ = 500.00, $\sigma_S^2$ = −408.33, and $\sigma_E^2$ = 183.33

Unconstrained OLS results in a negative estimation of variance

**b)** Set $\sigma_F^2$ to zero, OLS estimate using $S$ and $E$ and set estimate as: max{0, estimate}

| $y_i^{res} \times y_{i'}^{res}$ | *S* | *E* |
|---|---|---|
| 25 | 1 | 1 |
| 50 | 1 | 0 |
| 75 | 1 | 0 |
| 100 | 1 | 1 |
| 150 | 1 | 0 |
| 225 | 1 | 1 |
| 400 | 1 | 1 |
| 500 | 0 | 0 |
| 625 | 1 | 1 |

Estimated variance parameters:

$\sigma_F^2$ = 0.00
$\sigma_S^2$ = 91.67
$\sigma_E^2$ = 183.33

Predicted value = $\sigma_S^2 \times S + \sigma_E^2 \times E$

Cost = 489166.67

**c)** Set $\sigma_F^2$ and $\sigma_S^2$ to zero, OLS estimate using $E$ and set estimate as: max{0, estimate}

| $y_i^{res} \times y_{i'}^{res}$ | *E* |
|---|---|
| 25 | 1 |
| 50 | 0 |
| 75 | 0 |
| 100 | 1 |
| 150 | 0 |
| 225 | 1 |
| 400 | 1 |
| 500 | 0 |
| 625 | 1 |

Estimated variance parameters:

$\sigma_F^2$ = 0.00
$\sigma_S^2$ = 0.00
$\sigma_E^2$ = 275.00

Predicted value = $\sigma_E^2 \times E$

Cost = 514375.00

**d)**

| Configuration | Parameter set to zero | Estimated parameters: max{0,estimate} | | | Cost |
|---|---|---|---|---|---|
| | | $\sigma_F^2$ | $\sigma_S^2$ | $\sigma_E^2$ | |
| Configuration 1 | None | 500.00 | 0 | 183.33 | 1573055.56 |
| Configuration 2 | $\sigma_F^2$ | 0 | 91.67 | 183.33 | 489166.67 |
| Configuration 3 | $\sigma_S^2$ | 193.75 | 0 | 81.25 | 364218.75 |
| Configuration 4 | $\sigma_E^2$ | 500.00 | 0 | 0 | 992500.00 |
| Configuration 5 | $\sigma_F^2, \sigma_S^2$ | 0 | 0 | 275.00 | 514375.00 |
| Configuration 6 | $\sigma_F^2, \sigma_E^2$ | 0 | 206.25 | 0 | 552187.50 |
| Configuration 7 | $\sigma_S^2, \sigma_E^2$ | 238.89 | 0 | 0 | 378888.89 |
| Configuration 8 | $\sigma_F^2, \sigma_S^2, \sigma_E^2$ | 0 | 0 | 0 | 892500.00 |

**Figure S2:** Illustration of non-negative least squares estimation that is used for estimating the variance parameters for the random effects. **a)** Consider an example where there is family structure and repeated observations (two families with the first having three repeated measurements from a single subject and the second family having two subjects without any repeated measurements); the squares and products of $y^{res}$ are computed similar to **Figure S1**. Then, performing an ordinary least squares estimation without any constraints will result in one of the variance parameters $\sigma_S^2$ being negative (highlighted in orange color). In such a situation, we list all $2^z$ configurations (where, $z$ is the total number of random effects) of random effects ranging from considering all of them to dropping all of them. For each of these configurations, we re-estimate the variance parameters using ordinary least squares and take the maximum of zero and the estimate (i.e., negative estimates are set to zero). Additionally, we compute the "cost" associated with each estimate which is simply the sum of squared differences between the observed values ($y^{res} \times y^{res}$ column) and the predicted value (generically, the product of $\sigma$ and the design matrix). We show a worked-out example for two such scenarios in **b)** and **c)**. **d)** Once the parameters are estimated for all configurations and the associated "costs" computed, the set of parameter values with the minimum cost is selected as the solution for this estimation process (highlighted in green color).

#### 1.2 Implementation details

##### 1.2.1 Random effects

The current implementation of FEMA is designed to account for a wide range of practically relevant random effects. Specifically, users of FEMA can specify the following terms as random effects (independently, or in combination thereof): family effect ($F$, variance attributed to living in the same family), subject effect ($S$, variance attributed to repeated measurements), additive genetic effect ($A$, variance attributed to the additive genetic effect; this is specified as a $N \times N$ matrix with each entry corresponding to the genetic relationship between two observations – this is commonly calculated as the correlation coefficient between the standardized genotyping matrix and is referred as the genetic relationship matrix or GRM in the genetics field), dominant genetic effect ($D$, variance attributed to the dominant genetic effect; this is the square of the $A$ effect), maternal effect ($M$, variance attributed to having the same mother), paternal effect ($P$, variance attributed to having the same father), twin effect ($T$, variance attributed to being twins; this can be used to account for the fact that twins share the same fetal environment or for other kinds of factors like twins experiencing the same event at the same time at the same age, as opposed to siblings experiencing the same event in different forms due to the differences in their ages), and the home effect ($H$, variance attributed to sharing the same address or home). The random error term $E$, remains independent for each observation. A visual example of three different experimental designs is shown in Figure **Figure S3**. Aside from these random effects, it is straightforward to extend the FEMA code and implement any additional random effects of interest. An additional note with respect to the $A$ effect (and therefore the $D$ effect) is that it can be specified with theoretical expectations (for example, dizygotic twins, on average, have a genetic relationship of 0.5) or with empirical values (calculated as the correlation coefficient between the standardized genetic relationship matrix, where the standardization can be performed with theoretical allele frequencies or observed allele frequencies; see, for example, (Yang et al., 2011, 2010)).

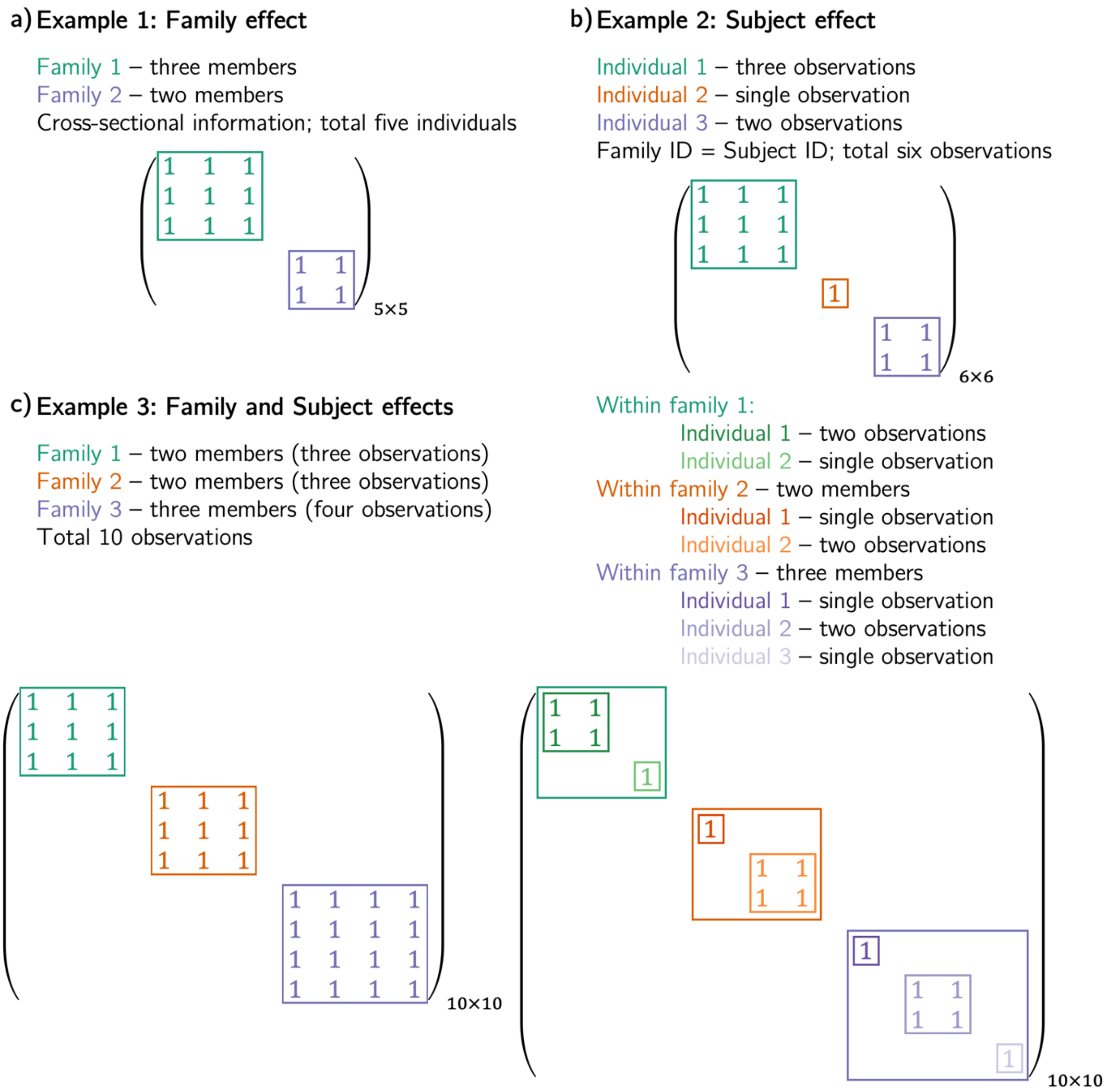


**d) Supported random effects in FEMA:**

| | |
|---|---|
| Family effect (*F*): | variance attributed to living in the same family |
| Subject effect (*S*): | variance attributed to repeated measurements |
| Additive genetic effect (*A*): | variance attributed to additive genetic effect |
| Dominant genetic effect (*D*): | variance attributed to dominant genetic effect |
| Maternal effect (*M*): | variance attributed to having the same mother |
| Paternal effect (*P*): | variance attributed to having the same father |
| Twin effect (*T*): | variance attributed to being twins (e.g., shared fetal environment) |
| Home effect (*H*): | variance attributed to sharing the same address or home |
| Error (*E*): | unexplained variance, independent for each observation (always included) |

**Figure S3:** Demonstration of how the design matrix for random effects ($ZZ^T$) is constructed within FEMA. **a)** a scenario with only family effect; **b)** a scenario with only repeated measurements; **c)** a scenario with both family design as well as repeated measurements; and **d)** list of random effects currently supported in FEMA.

##### 1.2.2 Sparsity of the random effects

A key detail in FEMA is the sparsity in the random effects design matrix, that we leverage for efficiency. For example, consider an $F$ and $S$ model where there are repeated measurements within a family, in addition to the independent observation-specific error terms ($E$). In such a situation, all calculations (for both the fixed effects and the random effects) are performed by only considering the individuals *within* each family. For example, when implementing equation (6), instead of computing squares and products of residuals across all observations (which would result in $(n \times (n+1))/2$ elements, we only compute squares and products of residuals within each family and omit the terms between families (see, also, **Figure S1**). We follow the same principle when considering other random effects, including the additive or dominant genetic effects. The same sparse formulation is also used when the user specifies ML as an estimator. This sparsity reduces memory requirement and helps in speeding up computation.

Once the random effects have been estimated, the fixed effects are re-estimated using generalized least squares (GLS; re-estimated because we start with an initial OLS solution for the fixed effects, compute the random effects, and then update the fixed effects again using GLS). Examining equations (3) and (7) (from the main text) in light of the sparsity of the random effects implies that the computation of the $V$ term is performed within families as well. Therefore, in the case of an $F$ and $S$ model (with an additional $E$ term), FEMA distinguishes between different family types based on the number of siblings in the family, and the presence/absence of longitudinal effect within the family. For all families that are of the same type, the $V$ term is calculated once and does not need be re-calculated again. As an example, consider the case that all families in a dataset have the same number of individuals and the same number of repeated observations per individual: in such a case, the covariance matrix $V$ will remain the same across all families and therefore only needs to be computed once.

##### 1.2.3 Binning strategy

Another important implementation detail to note is the use of a binning strategy during the estimation of the fixed effects. The key idea here is that if the random effects are similar to each other (across different imaging variables of the same type), then the same covariance parameters can be used across these variables for implementing the GLS solution for fixed effects. The $K$ bins, where $K \ll J$, are specified as a uniformly spaced multidimensional grid, where the spacing is based on a factor $1/K$. Therefore, each grid defines a range of values for the variance parameter of each random effect (see panel b) of **Figure S4**). Then, for all estimated variance parameters (estimated, for example, using MoM), we find the grid which covers that range of parameter values and assign the grid number as the bin number for that imaging variable (see panel c) of **Figure S4**). Each bin, in turn, specifies a set of variance components that multiply into the design matrices to give a total covariance matrix which is used in GLS to estimate the fixed effects. The granularity of the bins can be specified by the user as a single number (bin value) and is used to control the granularity of the estimation of the fixed effects – a bin value of one would mean that all $J$ imaging measures will have the same random effects covariance matrix $V$. We emphasize that the binning of the imaging variables is performed based on the estimated variance parameters of the random effects – i.e., binning is not based on *a priori* assumptions about the random effects but rather based on the estimates. This implies that voxels/vertices (or any imaging variable) that are not spatially contiguous may be binned together simply because they have similar estimated variance parameters. This binning allows us to efficiently implement the GLS solution for the fixed effects: $\hat{\beta}_j = \left(X^T V_j^{-1} X\right)^{-1} X^T V_j^{-1} y_j$; specifically, the binning strategy allows us to reuse the total covariance matrix $V$ across all imaging variables that are in the same bin. In our simulations, we have demonstrated that a finite number of gridded bins is sufficient to capture the variance components for all input imaging measures; specifically, we have shown that using a bin value of 20 dramatically reduces the required computational time with negligible negative effect on the accuracy of the estimates.

a) Estimate variance parameters of random effects for each variable

Assume nine imaging variables with these estimated variance parameters $V_F$, $V_S$, and $V_E$

| Variable 1 | Variable 2 | Variable 3 |
|---|---|---|
| 0.12 | 0.17 | 0.09 |
| 0.40 | 0.72 | 0.69 |
| 0.48 | 0.12 | 0.23 |
| **Variable 4** | **Variable 5** | **Variable 6** |
| 0.70 | 0.12 | 0.29 |
| 0.23 | 0.75 | 0.34 |
| 0.07 | 0.13 | 0.37 |
| **Variable 7** | **Variable 8** | **Variable 9** |
| 0.25 | 0.28 | 0.35 |
| 0.31 | 0.42 | 0.31 |
| 0.45 | 0.31 | 0.34 |

b) Define uniformly spaced grid of random effects (remove values where $V_F + V_S \neq 1$)

Bin size = 5, spacing = 1/5 = 0.20

| Bin # | $V_F$ | | $V_S$ | |
|---|---|---|---|---|
| | Lower | Upper | Lower | Upper |
| 1 | 0.00 | 0.20 | 0.00 | 0.20 |
| 2 | 0.20 | 0.40 | 0.00 | 0.20 |
| 3 | 0.40 | 0.60 | 0.00 | 0.20 |
| 4 | 0.60 | 0.80 | 0.00 | 0.20 |
| 5 | 0.80 | 1.00 | 0.00 | 0.20 |
| 6 | 0.00 | 0.20 | 0.20 | 0.40 |
| : | : | : | : | : |
| 19 | 0.20 | 0.40 | 0.80 | 1.00 |

Bin size = 20, spacing = 1/20 = 0.05

| Bin # | $V_F$ | | $V_S$ | |
|---|---|---|---|---|
| | Lower | Upper | Lower | Upper |
| 1 | 0.00 | 0.05 | 0.00 | 0.05 |
| 2 | 0.05 | 0.10 | 0.00 | 0.05 |
| 3 | 0.10 | 0.15 | 0.00 | 0.05 |
| 4 | 0.15 | 0.20 | 0.00 | 0.05 |
| 5 | 0.25 | 0.30 | 0.00 | 0.05 |
| 6 | 0.30 | 0.35 | 0.00 | 0.05 |
| : | : | : | : | : |
| 229 | 0.05 | 0.10 | 0.95 | 1.00 |

c) Assign bins based on similarity of grid values and estimated variance values

| Variable 1 | Variable 2 | Variable 3 |
|---|---|---|
| Bin 11 | Bin 15 | Bin 15 |
| 0.00 - 0.20 | 0.00 - 0.20 | 0.00 - 0.20 |
| 0.40 - 0.60 | 0.60 - 0.80 | 0.60 - 0.80 |
| **Variable 4** | **Variable 5** | **Variable 6** |
| Bin 9 | Bin 15 | Bin 7 |
| 0.60 - 0.80 | 0.00 - 0.20 | 0.20 - 0.40 |
| 0.20 - 0.40 | 0.60 - 0.80 | 0.20 - 0.40 |
| **Variable 7** | **Variable 8** | **Variable 9** |
| Bin 7 | Bin 12 | Bin 7 |
| 0.20 - 0.40 | 0.20 - 0.40 | 0.20 - 0.40 |
| 0.20 - 0.40 | 0.40 - 0.60 | 0.20 - 0.40 |

| Variable 1 | Variable 2 | Variable 3 |
|---|---|---|
| Bin 142 | Bin 206 | Bin 196 |
| 0.00 - 0.15 | 0.15 - 0.20 | 0.05 - 0.10 |
| 0.40 - 0.45 | 0.70 - 0.75 | 0.65 - 0.70 |
| **Variable 4** | **Variable 5** | **Variable 6** |
| Bin 92 | Bin 212 | Bin 116 |
| 0.70 - 0.75 | 0.10 - 0.15 | 0.25 - 0.30 |
| 0.20 - 0.25 | 0.75 - 0.80 | 0.30 - 0.35 |
| **Variable 7** | **Variable 8** | **Variable 9** |
| Bin 100 | Bin 145 | Bin 117 |
| 0.25 - 0.30 | 0.25 - 0.30 | 0.30 - 0.35 |
| 0.25 - 0.30 | 0.40 - 0.45 | 0.30 - 0.35 |

**Figure S4**: Demonstration of the FEMA binning strategy. **A)** Assume a case of nine imaging variables with different estimates of variance parameters for family (green color) and subject effects (purple color; and additionally, unmodeled error term shown in orange color). **B)** Based on the user-specified bin value $K$, we create uniformly spaced bins where the grid spacing is based on $1/K$ – for example, a bin value of 5 would imply grid spacing by 0.20 while a bin value of 20 would imply a grid spacing by 0.05. **c)** Based on the estimated variance parameters for each random effect, each imaging variable is assigned a bin value; for example, variable 2 has $V_F = 0.17$ and $V_S = 0.72$ and has been assigned to bin 15 (when bin size is 5) – this is because bin number 15 covers the range of variance parameters $V_F = 0.00 - 0.20$ and $V_S = 0.60 - 0.80$. On the other hand, when using a bin value of 20, the same variable is assigned to bin number 206 (which covers the range $V_F = 0.15 - 0.20$ and $V_S = 0.70 - 0.75$). As can be seen from the left side of panel **c)**, variables 2, 3, and 5 have all been assigned to bin 15 while variables 6, 7, and 9 are assigned to bin 7; other variables belong to other bins. On the other hand, on the right side of panel **c)**, each variable was assigned to a separate bin as the estimated variance parameters of the random effects were not similar enough. For all imaging variables which are assigned the same bins, we implement the generalized least squares (GLS) solution for the fixed effects in one go, thereby dramatically bringing down computational time.

###### 1.2.4 Using maximum likelihood estimator

In addition to using method of moments (MoM) for estimating variance components, users also have the option of selecting a ML estimator, instead of the MoM estimator. The ML estimator works by applying a minimization of the negative log likelihood of the multivariate normal probability density function, with non-negativity constraint on the estimates of the variance parameters. If the user requests ML estimates of the parameters, the starting point for the constrained minimization is based on the MoM solution, thereby allowing us to converge to the ML solution faster (since the MoM solution will be closer to the final solution). However, we note that unlike the MoM formulation, it is not straightforward to apply ML to $J$ imaging variables in a single computation; therefore, in this case, we loop over every imaging measure in $J$, which can be quite time consuming for large values of $J$. We additionally note here that REML estimators are the default choice in some implementations that solve mixed models (for example, in the *lmer* package in R (Bates et al., 2015)). The ML estimation of random effects does not account for the loss in degrees of freedom due to estimating the fixed effects (see, for example, (Harville, 1977; Searle et al., 1992, pp. 249–250)), thereby resulting in biased estimates. However, this bias is small for large samples, which is primarily the use case for FEMA. Therefore, in our implementation, we provide an optional use of ML as the estimator.

###### 1.2.5 Uncertainty estimation

Within FEMA, we provide two ways of calculating the confidence intervals on the variance parameters of the random effects – one using a wild bootstrap method and another by using profile likelihood. Within the wild bootstrap, for every bootstrap iteration $b$, we resample the residuals obtained after estimating the fixed effects using GLS (i.e., after implementing equation (7) from the main text). After the first calculation of the random effects and the fixed effects, the predicted outcome variable is $\hat{y} = X\hat{\beta}_{GLS}$ and the marginal residuals are $y^{res} = y - \hat{y}$. Then, using these values, the bootstrap data generation process is specified as:

$$y_b = \hat{y} + wy^{res} \quad (4)$$

where, $w$ is a random variable sampled from a standard normal distribution. Using the new $y_b$, we re-estimate the variance components and re-implement the GLS solution to get new estimates of fixed effects. We repeat this process $\mathcal{B}$ times (where $\mathcal{B}$ is specified by the user). Then, the 95% confidence intervals for the random effects can be calculated as the 2.5$^{th}$ and 97.5$^{th}$ percentiles – i.e., the values at $0.025 \times \mathcal{B}$ and $0.975 \times \mathcal{B}$ are the lower and upper limits of the 95% bootstrap confidence intervals. A similar method can be used for calculating the bootstrap confidence intervals on fixed effects (although one could use the estimated coefficient and standard error for computing the confidence intervals). Similar to other operations, we perform the wild bootstrap resampling of the outcome variable for each family or cluster in the data (specifically, $w$ is randomly drawn from a standard normal distribution separately for each family or cluster). For calculating the confidence intervals using a profile likelihood, the user must specify the random effects estimator as ML. In this case, each random effect is considered one at a time (while the other random effects have fixed values) and the likelihood is calculated within a range of parameter values. The details of computing this partially maximized log likelihood (or profile log likelihood) can be found in (Cole et al., 2014; Fitzmaurice et al., 2011, p. 99; McCullagh and Nelder, 1989, pp. 254–255; Murphy and van der Vaart, 2000).

##### 1.2.6 Number of iterations

Another point to note is that within FEMA, users have an option of specifying the number of iterations (default is one). If the number of iterations is set to be greater than one, then the estimation of the variance of the random effects and the slope for the fixed effects is repeated as many times as the number of iterations. This can, theoretically speaking, improve the estimation accuracy of the coefficients. However, in practice, we have only seen minor changes in the estimates with higher number of iterations. If the number of iterations is specified as greater than one, and the user enables permutation testing for generation of wild bootstrap-based confidence intervals, each permutation would involve performing of the set number of iterations.

##### 1.2.7 Reporting of variance components

Another implementation detail that we note about FEMA is that the variance parameters are reported in units of proportion of residual variance in the data. To clarify, once the fixed effects are regressed from the $y$ variables, the marginal residual is the total residual variance in the data. Then, the estimated variance parameters are converted as proportions of this total residual variance in the data. The same normalization is applied for the (wild bootstrap) permutated variance parameters, which are expressed as proportions of permuted total residual variance in the data. The rationale for this scaling is that it directly connects the variance of the additive genetic effect, $A$, to the notion of (genetic) heritability (i.e., the proportion of variance in the $y$ variable attributable to the genetic variation). Additionally, it allows a comparison of the variances across different $y$ variables, irrespective of the differences in their total residual variances. Since the total residual variance in the data is also saved as an output, the user has an option of converting the variances back to the original scale.

###### 1.2.8 Parallel computing

While FEMA is computationally efficient, providing estimates for a large number of imaging variables in a short time (see results of simulations 3), it is possible to further speed up computation by using parallel computing. Specifically, the FEMA code is written in a way that the "family structure" can be re-used. In other words, the parsed random effects, which are used to create the covariance matrix for the estimation of variance parameters for the random effects, can be re-used across all imaging variables. This allows us to split the computation across the bins, where each bin is a separate "job" while sharing the same family structure. Thus, it is possible to speed up FEMA computation by several folds, thereby allowing processing of large-scale whole-brain imaging data in almost real time (especially if the parallel processing workers are persistent workers, thereby eliminating overhead costs of starting a new worker).

### 2. Results

#### 2.1 Simulation results

##### *2.1.1* Comparison with *fitlme*

###### *2.1.1.1 Fixed effects*

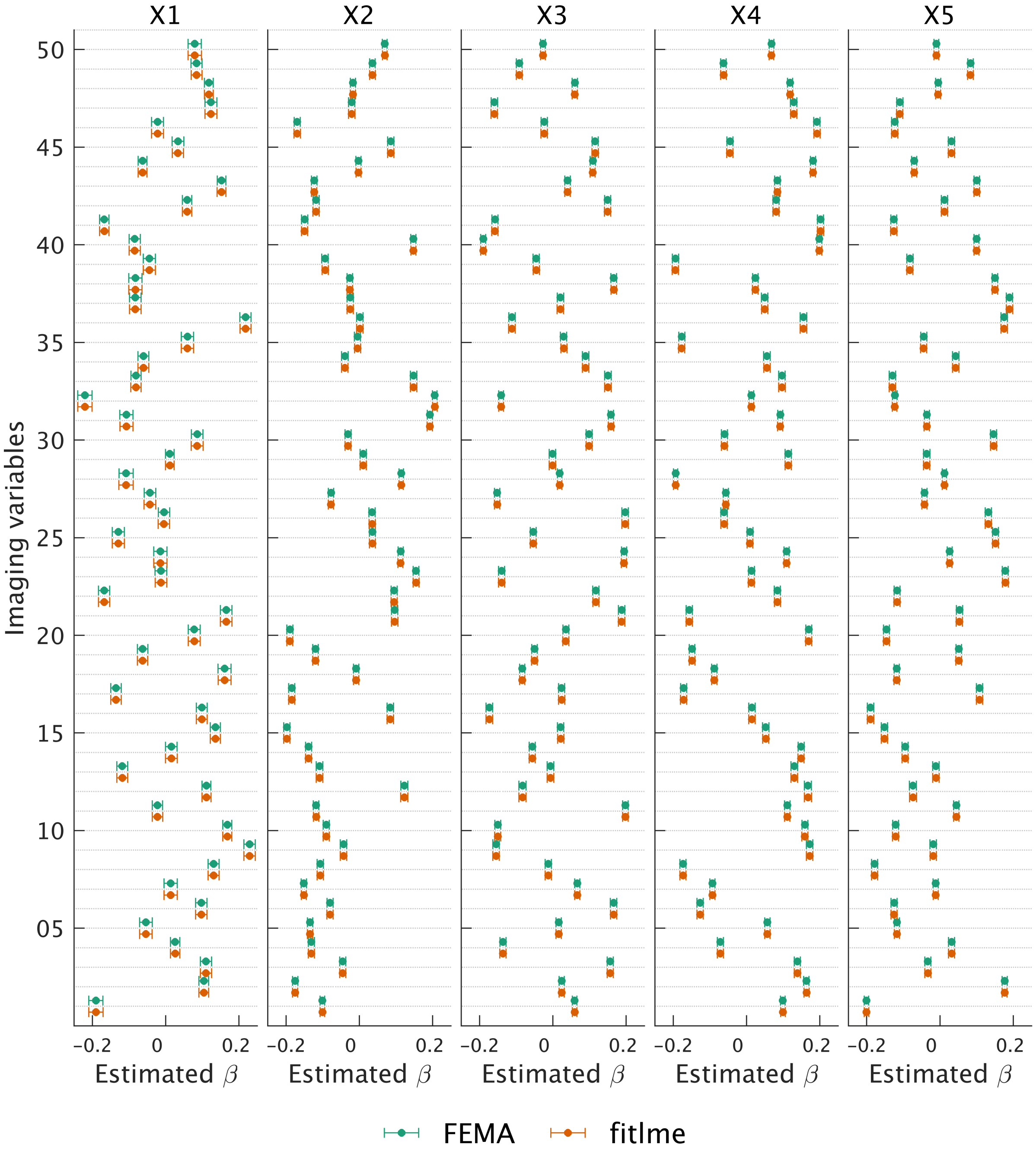


**Figure S5:** Comparison of FEMA estimated slopes (green points) for the five fixed effects ($X1$ to $X5$) across 50 imaging variables with the slopes estimated from MATLAB's *fitlme* function (orange points). The error bars around the point estimates represent the standard error.

*2.1.1.2 Random effects*

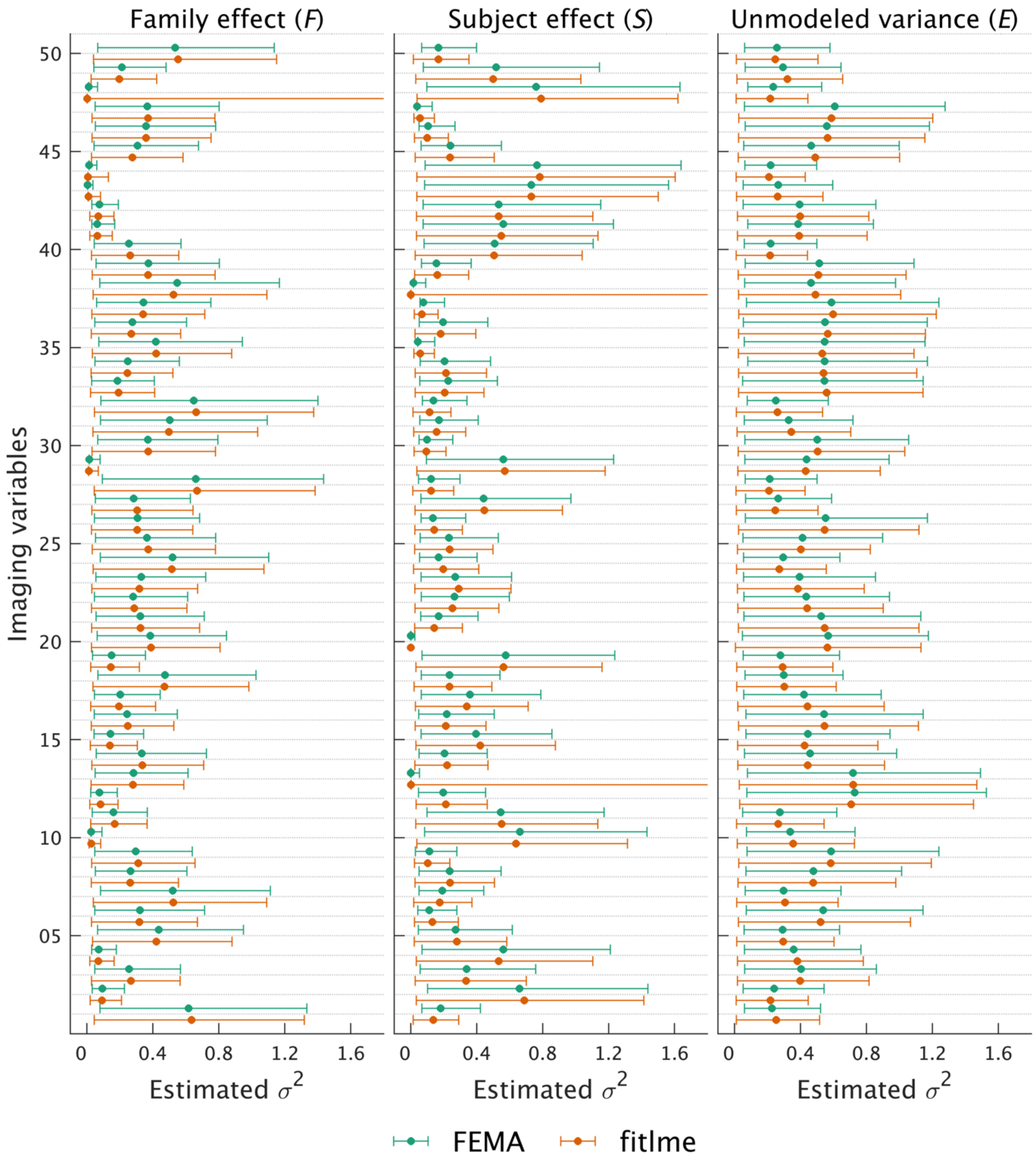


**Figure S6:** Comparison of the variance parameter estimates of the three random effects (family effect $F$, subject effect $S$, and the unmodeled variance term $E$) from FEMA (green points) and MATLAB's *fitlme* (orange points). The error bars around the point estimates are the 95% confidence intervals (estimated in FEMA by running 100 wild bootstrap permutations). Note that for one imaging variable for the family effect, and two imaging variables for the subject effect, MATLAB's confidence interval upper limit were unreasonably large and are truncated in this figure. For one imaging variable, for the subject effect, MATLAB returned a NaN (not a number) value for the confidence interval. Additionally, note that the lower limits of the confidence intervals are truncated at zero because variance cannot be negative.

##### 2.1.2 Computational time as a function of number of observations

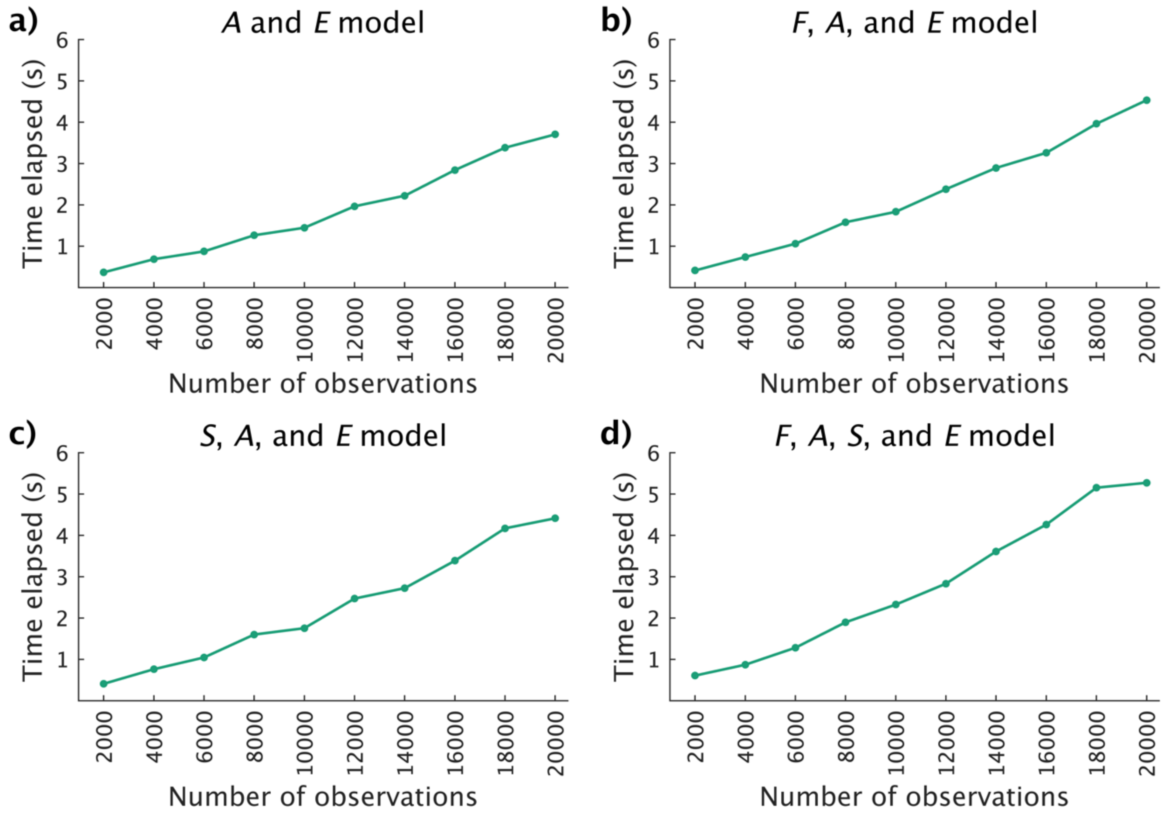


**Figure S7:** Time taken by FEMA to fit different models as a function of increasing number of observations for 50 imaging variables and five fixed effects. The random effects included in each model are indicated at the top of each panel: family effect $F$, additive genetic effect $A$, subject effect $S$, and the unmodeled variance term $E$.

##### 2.1.3 Type I error rate

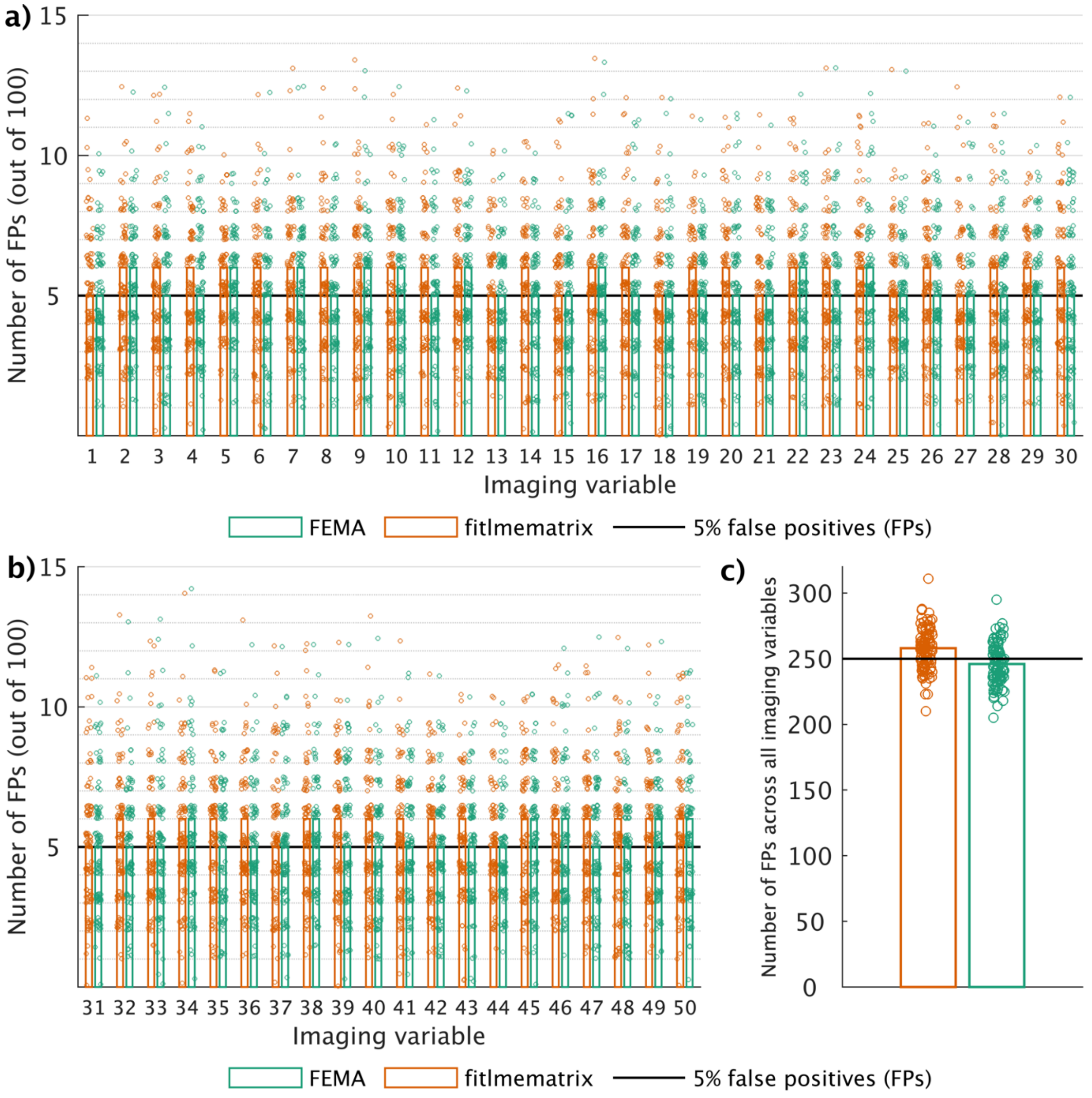


**Figure S8:** Comparison of type I error rate for fixed effects estimation between MATLAB's *fitlmematrix* function and FEMA; we performed 100 repetitions of simulating 10,000 observations and 50 imaging variables, with each imaging variable associated with 100 $X$ variables (zero effect) and family and subject effects specified as random effects; panel **a)** shows the number of false positives (FPs) across the 100 fixed effects for the first 30 imaging variables (i.e., for every imaging variable, we counted the number of $X$ variables which had a $p$ value less than 0.05) – the dots represent the number of FPs for every repeat while the bars show the average number of FPs across 100 repeats rounded towards positive infinity; panel **b)** shows the number of FPs across the 100 fixed effects for the remaining 20 imaging variables; and panel **c)** shows the total of number of FPs across all 50 imaging variables with the dots representing the sum (across 50 imaging variables) of FPs for every repetition while the bars represent the average number of FPs rounded towards positive infinity; the solid black line represents the 5% FP rate. Note that the data points are jittered to allow for greater visibility.

###### 2.1.4 Effect of number of observations

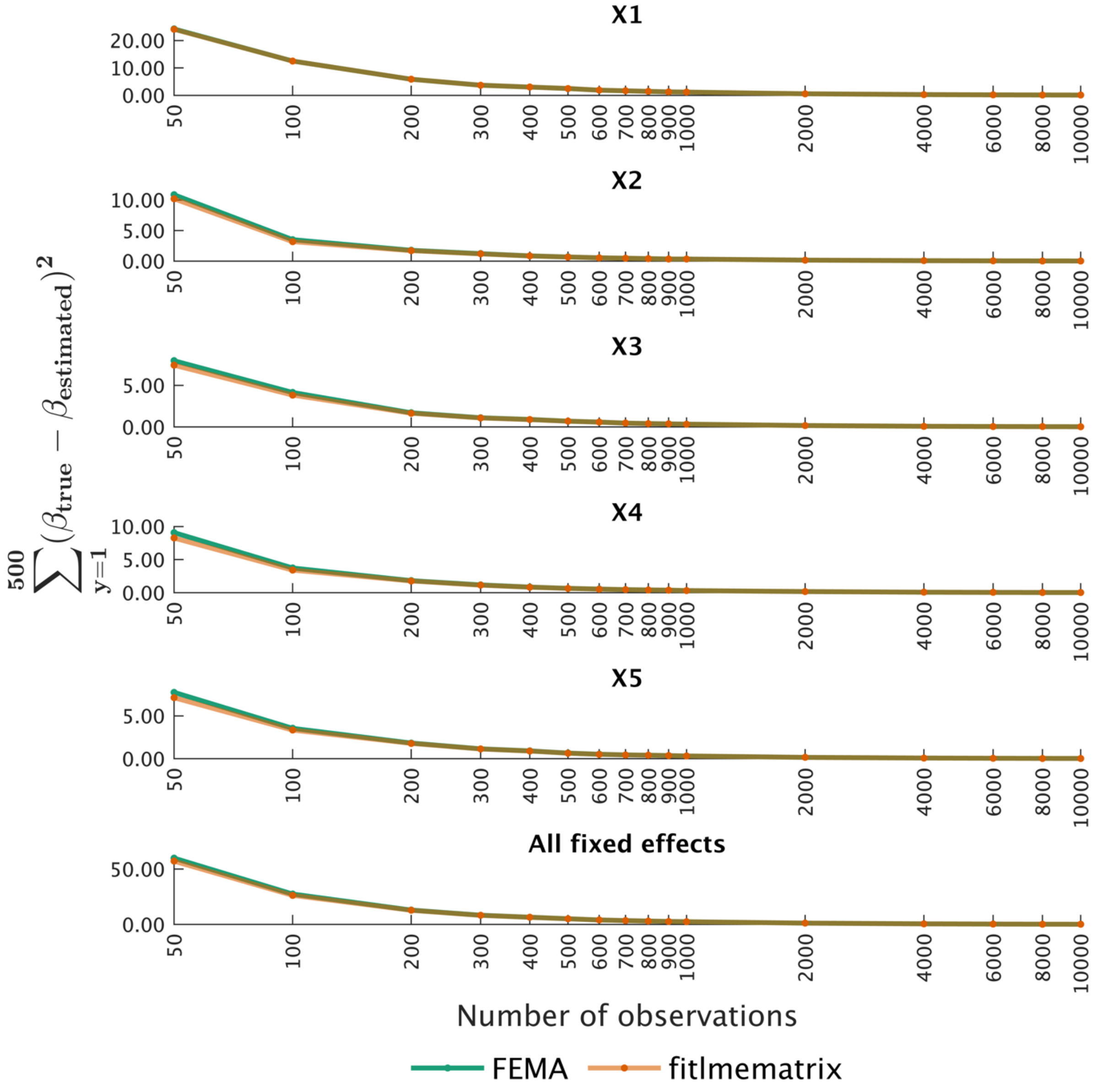


**Figure S9:** Comparison of ground truth and point estimates for fixed effects from FEMA (at a bin value of 20; green) and MATLAB's *fitlmematrix* (orange) as a function of increasing number of observations. We simulated five repeats of 500 outcome variables, five fixed effects, and random effects of family and subject, at different sample sizes (number of observations); each simulation had up to five family members and up to five repeated observations. For every sample size, for every outcome variable, for every fixed effect, we calculated the squared difference between the ground truth and the estimated parameters from FEMA and *fitlmematrix*; then, we averaged this squared difference across the five repeats. Next, we summed the average squared difference across the 500 outcome variables. Finally, we summed these across the five fixed effects to get a single estimate of the difference between ground truth and the parameter estimates from FEMA and *fitlmematrix*. Note that the *x*-axis is non-linear.

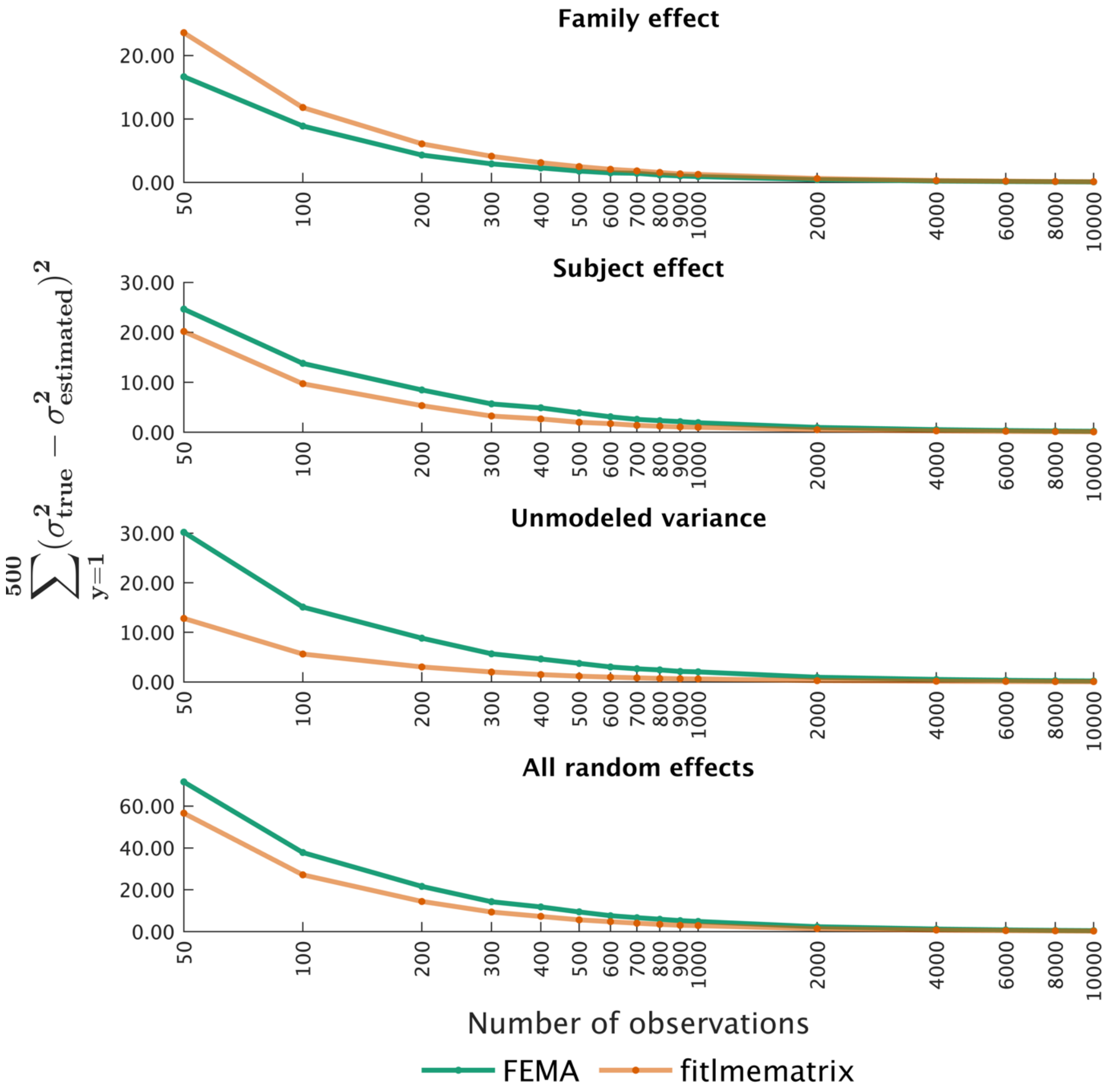


**Figure S10:** Comparison of ground truth and point estimates for random effects from FEMA (at a bin value of 20; green) and MATLAB's *fitlmematrix* (orange) as a function of increasing number of observations. We simulated five repeats of 500 outcome variables, five fixed effects, and random effects of family and subject, at different sample sizes (number of observations); each simulation had up to five family members and up to five repeated observations. For every sample size, for every outcome variable, for every random effect, we calculated the squared difference between the ground truth and the estimated parameters from FEMA and *fitlmematrix*; then, we averaged this squared difference across the five repeats. Next, we summed the average squared difference across the 500 outcome variables. Finally, we summed these across the three random effects to get a single estimate of the difference between ground truth and the parameter estimates from FEMA and *fitlmematrix*. Note that the *x*-axis is non-linear.

##### 2.1.5 Type I error rate as a function of bin values

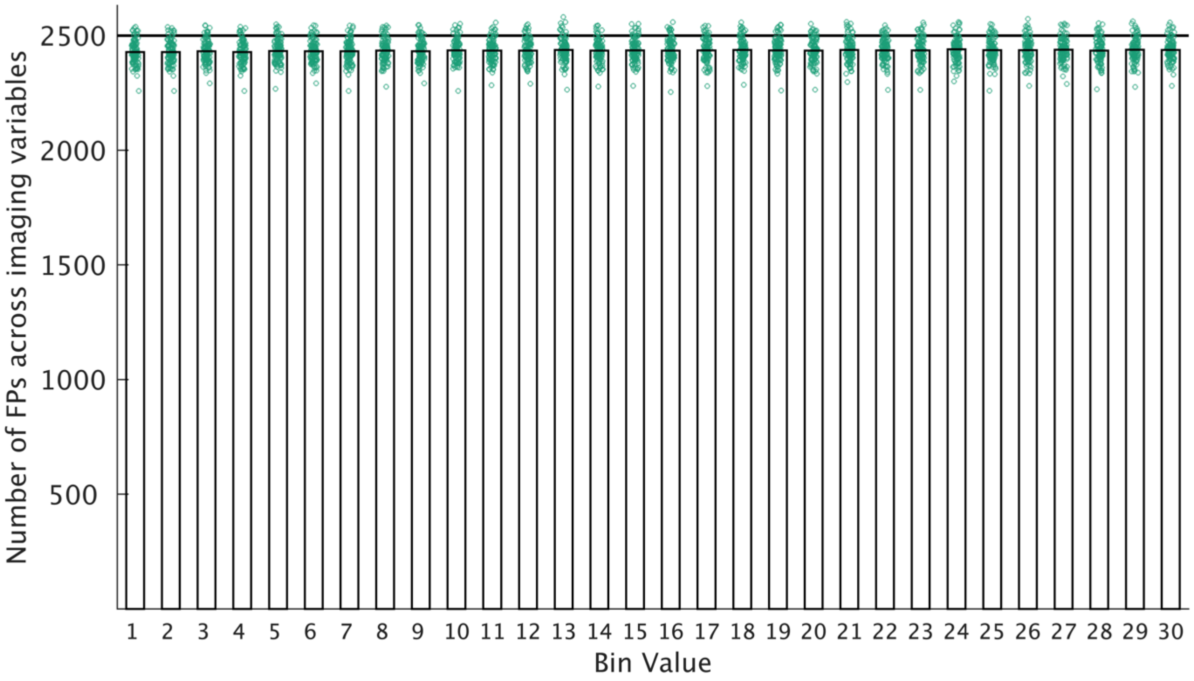


**Figure S11:** Type I error rate as a function of bin values for fixed effects estimation; we performed 100 repetitions of simulating 10,000 observations and 500 imaging variables, with each imaging variable associated with 100 $X$ variables (zero effect) and family and subject effects specified as random effects; for every repeat, for every bin value, we counted the number of false positives (i.e., the number of $X$ variables which had a $p$ value less than 0.05); the green dots indicate the number of false positives (FPs) at each bin value across the imaging variables (for each repeat) while the bars show the average (across repeats) number of FPs for that bin size (rounded to positive infinity); the solid black line represents the 5% FP rate. Note that the data points are jittered to allow for greater visibility.

#### 2.2 Empirical results

##### 2.2.1 ROI-level cortical thickness

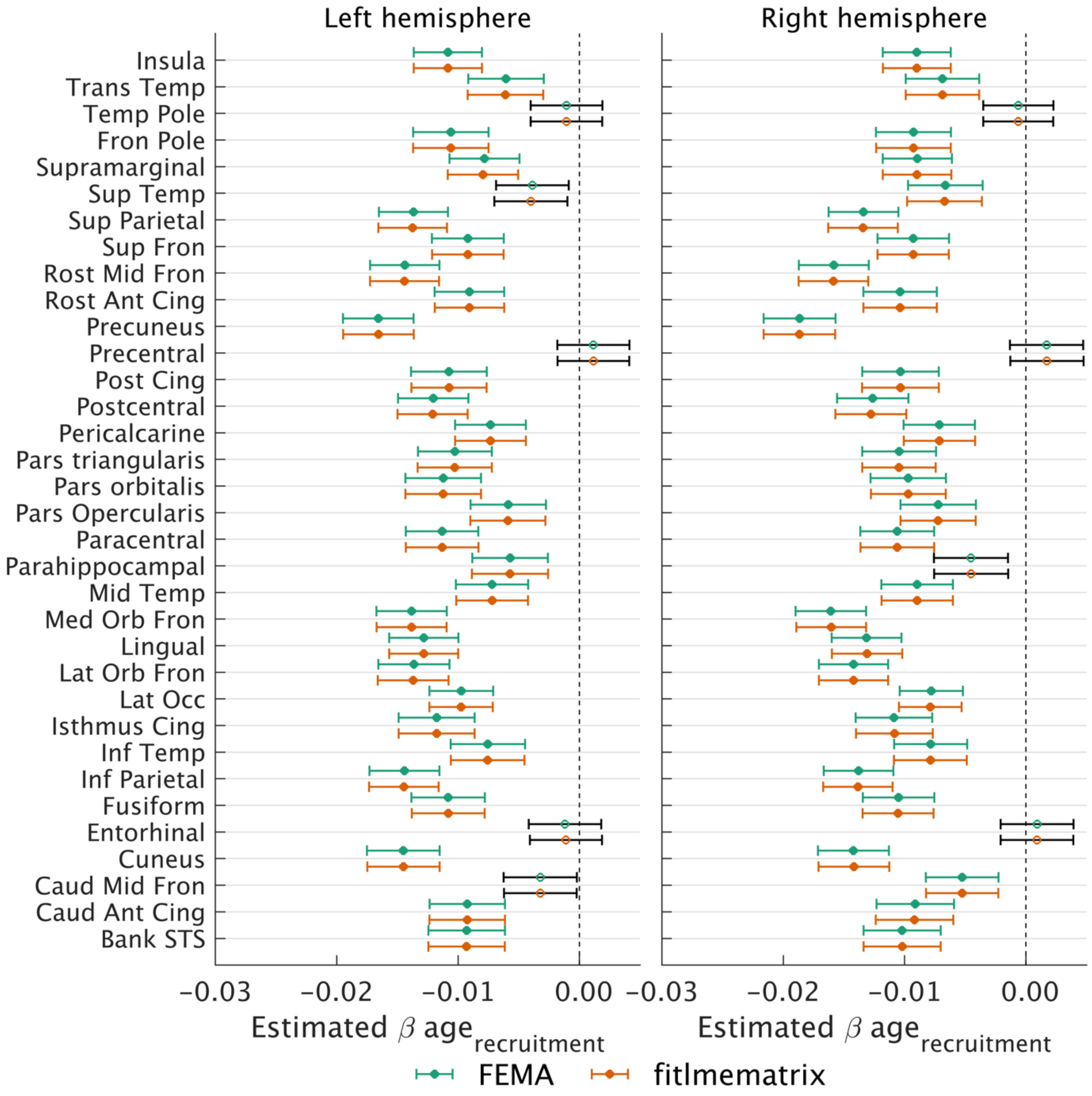


**Figure S12:** Comparison of the estimated slopes and 95% confidence intervals (calculated as $1.96 \times SE$) for the cross-sectional effect of age, $age_{recruitmemt}$, between MATLAB's *fitlmematrix* (orange points) and FEMA (at a bin value of 20, green points). The coefficients which were statistically significant (after a Bonferroni correction for 68 ROIs) are indicated with filled points and colored confidence interval lines while the estimates for the regions which did not survive multiple comparison correction are marked with non-filled in points and with black lines for their confidence intervals; Ant = anterior; Caud = caudal; Cing = cingulate; Fron = frontal; Inf = inferior; Lat = lateral; Med = medial; Mid = middle; Occ = occipital; Orb = orbital; Post = posterior; Rost = rotstral; STS = superior temporal sulcus; Sup = superior; Temp = temporal; Trans = transverse.

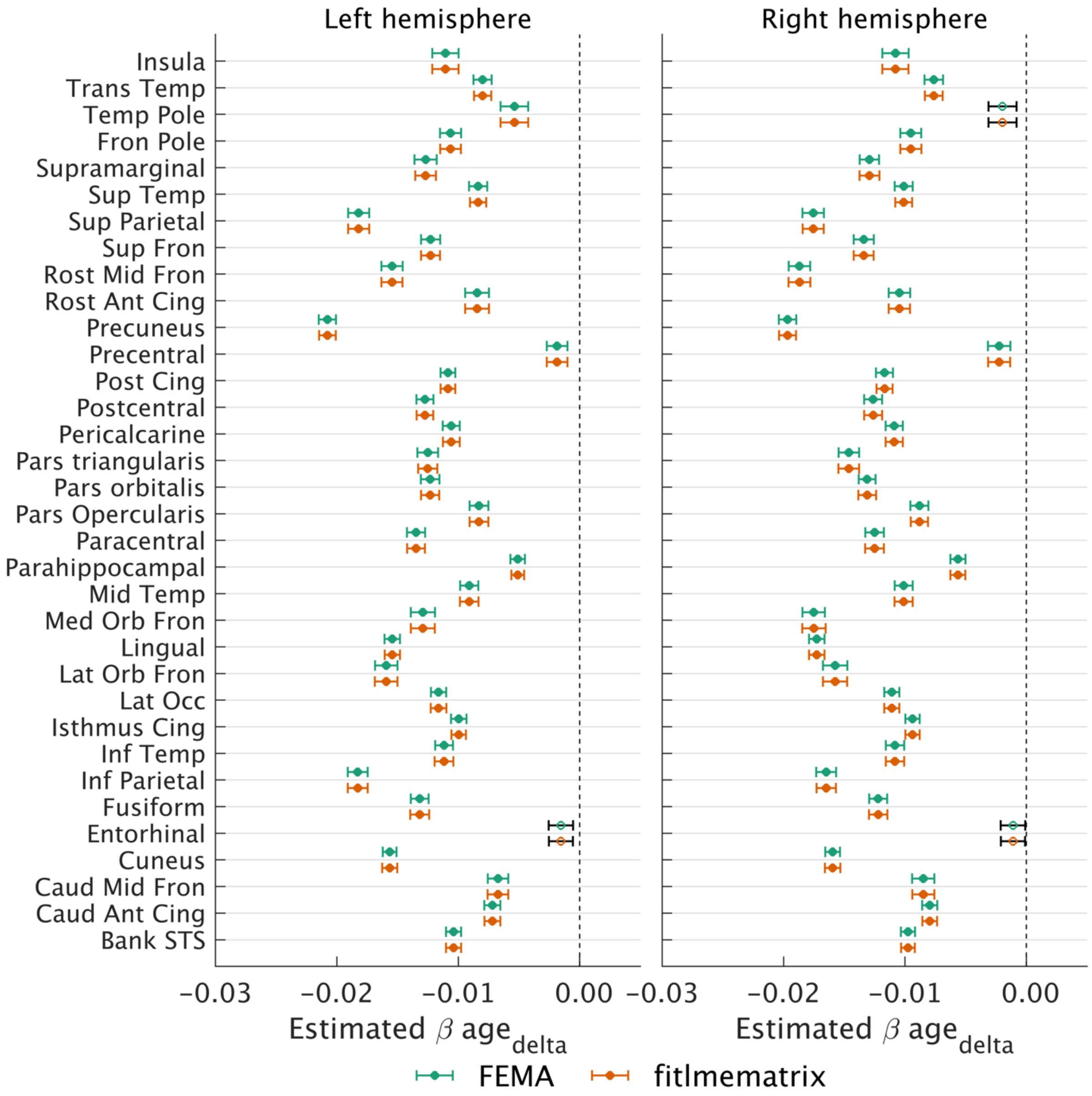


**Figure S13:** Comparison of the estimated slopes and 95% confidence intervals (calculated as $1.96 \times SE$) for the longitudinal effect of age, $age_{delta}$, between MATLAB's *fitlmematrix* (orange points) and FEMA (at a bin value of 20, green points). The coefficients which were statistically significant (after a Bonferroni correction for 68 ROIs) are indicated with filled points and colored confidence interval lines while the estimates for the regions which did not survive multiple comparison correction are marked with non-filled in points and with black lines for their confidence intervals; Ant = anterior; Caud = caudal; Cing = cingulate; Fron = frontal; Inf = inferior; Lat = lateral; Med = medial; Mid = middle; Occ = occipital; Orb = orbital; Post = posterior; Rost = rotstral; STS = superior temporal sulcus; Sup = superior; Temp = temporal; Trans = transverse.

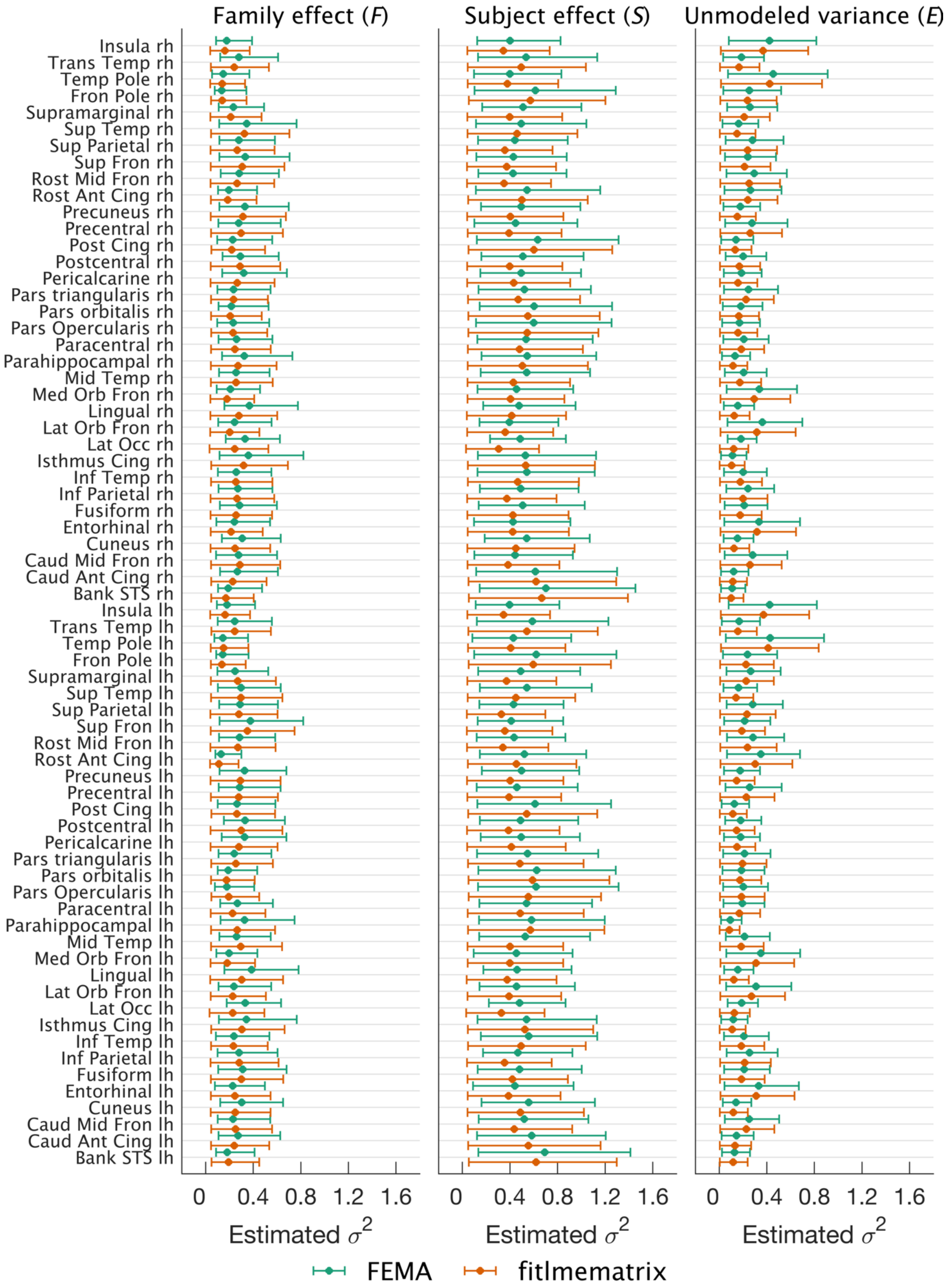


**Figure S14**: Comparison of estimates of variance of random effects and their confidence intervals for family effect ($F$), subject effect ($S$), and unmodeled variance ($E$) across cortical thickness from 68 regions of interest between MATLAB's *fitlmematrix* (orange points) and FEMA (at a bin value of 20, green points); Ant = anterior; Caud = caudal; Cing = cingulate; Fron = frontal; Inf = inferior; Lat = lateral; Med = medial; Mid = middle; Occ

= occipital; Orb = orbital; Post = posterior; Rost = rotstral; STS = superior temporal sulcus; Sup = superior; Temp = temporal; Trans = transverse; "lh" suffix corresponds to the left hemisphere and "rh" suffix to the ROI name corresponds to the right hemisphere.

##### 2.2.2 Vertex-wise cortical thickness

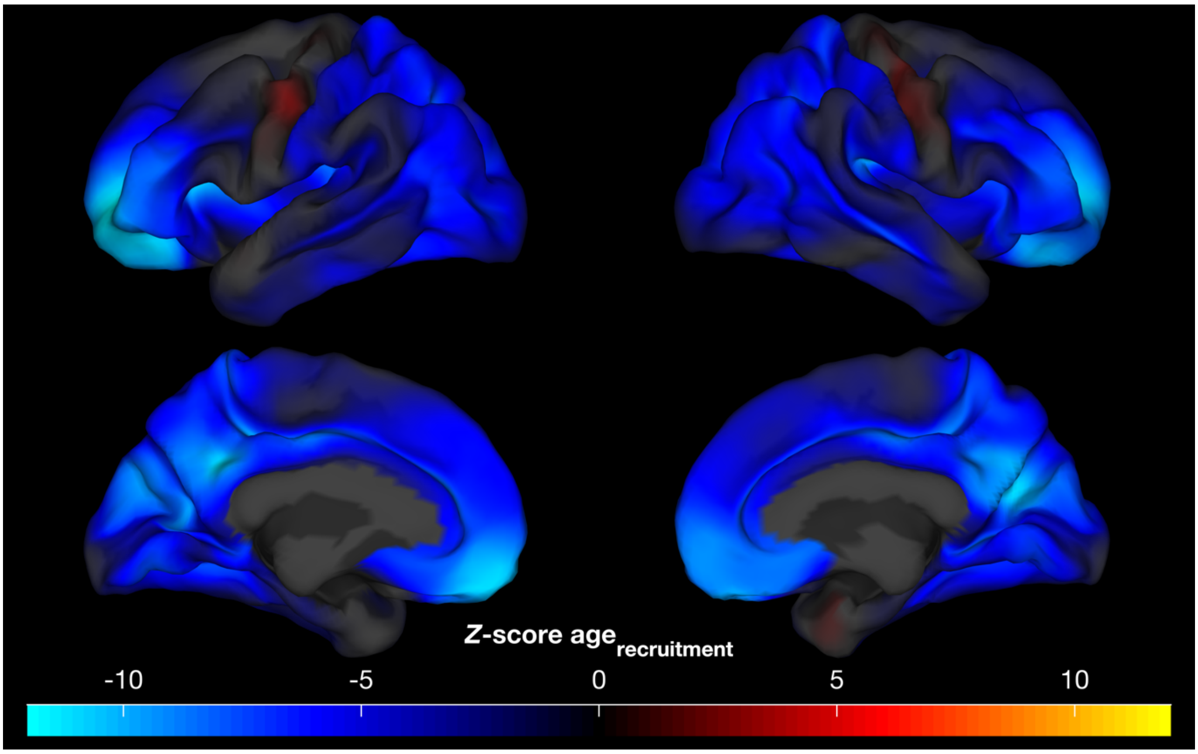


**Figure S15:** Unthresholded vertex-wise *Z* scores for the cross-sectional effect of age ($age_{recruitment}$); performing whole-brain vertex-wise analysis using FEMA took about 11 seconds.

##### 2.2.3 Connectome-wide functional connectivity

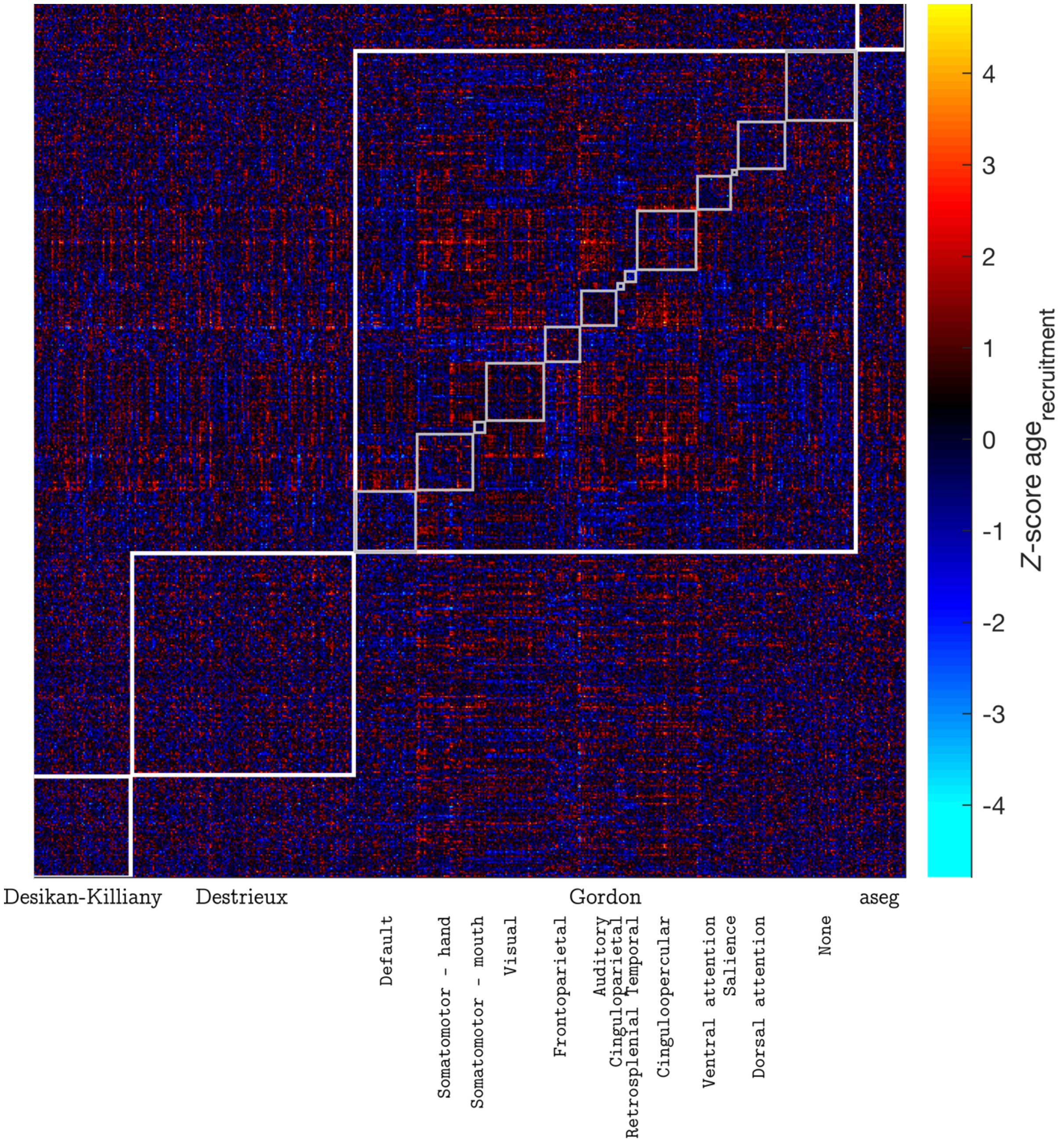


**Figure S16:** Distribution of *Z*-scores for the cross-sectional effect of age ($age_{recruitment}$) across 582 ROIs; the ROIs were defined using a combination of Desikan-Killiany atlas, Destrieux atlas, Gordon parcellation, and the aseg atlas; the Gordon parcellation is further divided into 13 communities; performing connectome-wide analysis using FEMA took about 54 seconds.
